# Single-cell and spatial transcriptomics uncover the role of B chromosomes in driving plant invasiveness

**DOI:** 10.1101/2024.12.31.630906

**Authors:** Cui Wang, James Ord, Mengxiao Yan, Yunfei Cai, Hongjin Shao, Lele Lin, Jarkko Salojärvi, Lele Liu, Weihua Guo

## Abstract

Invasive plants pose a major threat to global biodiversity, yet the molecular and genomic mechanisms underlying their success remain poorly understood. Here, we investigate the common reed (*Phragmites australis*), a grass species that became invasive in North America after introduction from Europe, to unravel the molecular mechanisms of its invasiveness. By integrating single-cell RNA sequencing, spatial transcriptomics, and comparative genomics, we constructed a single-cell atlas of *P. australis* and identified 19 transcriptionally distinct cell types, including shoot apical meristem, epidermal, and vascular tissues. Comparative analysis of common garden-grown native (European) and invasive (North American) populations revealed a significant proportion of differentially expressed genes (DEGs) located on B chromosomes, which underwent copy number expansion in invasive genomes. The proportion of B chromosome genes in DEGs varies across cell types, with the highest proportions observed in the epidermis and mesophyll, and lower proportions in the vascular tissues. Gene IMPA-3, a B chromosome gene likely derived from transposable element activity, exhibited an elevated mutation rate compared to its ancestral counterparts, potentially enhancing adaptive evolution in invasive populations. Invasive individuals also displayed molecular regulatory networks related to photosynthetic efficiency, stress tolerance, and growth-defense trade-offs. Together, our findings provide a cell-type-resolved molecular atlas of a non-model invasive plant and offer key insights into the cellular and genomic architecture of plant invasiveness, with implications for ecological management.

## Introduction

Biological invasions pose a profound threat to global ecosystems, as non-native species can disrupt ecological equilibrium, outcompete indigenous flora and fauna, and reshape habitat structures (*1*). While many species become invasive upon introduction to new environments, their counterparts may remain non-invasive in their native habitats, as seen in *Spartina* spp., *Ambrosia artemisiifolia* L., and *Phragmites australis* (*2, 3*). Invasive populations often exhibit distinct phenotypic or reproductive traits in the region they have invaded. These contrasting traits, such as altered growth rates, reproductive strategies, or resource use, can play a pivotal role in their invasive success(*4, 5*). Genetic mechanisms driving such divergence include copy number variation, epistasis, hybridization, and large-effect loci, as well as rapid genomic shifts involving transposable elements and pleiotropy (*6, 7*). Understanding these genetic mechanisms is essential for uncovering how species become invasive and how they can outcompete native species in newly occupied habitats(*8*). Traditionally, large-scale ecological observations or phylogenetic studies that use a limited number of genetic markers were used to infer the origins and dynamics of invasive species. With the growing accessibility of sequencing technologies, omics data has emerged as a powerful tool in ecological genetic research for invasive species (*9*).

The common reed (*Phragmites australis*) is one of the most globally widespread plant species, thriving in a variety of wetland habitats across temperate and tropical regions. Its extensive distribution is accompanied by remarkable intraspecific genetic diversity, with significant variation in genetic loci, ploidy levels, and morphological traits (*10–12*). This diversity has allowed *P. australis* to adapt to a wide range of environmental conditions, contributing to its ecological resilience and competitive success. Despite its native status in Europe, where it plays an essential role in wetland ecosystems without exhibiting invasive tendencies, *P. australis* has become a highly aggressive invasive species in North America. Its introduction to the continent is believed to have occurred approximately 200 years ago(*3*). Since then, *P. australis* has expanded rapidly across North American wetlands, where it outcompetes native vegetation, reduces biodiversity, and alters hydrological and nutrient cycling processes(*13*).

One of the key differences observed between invasive and native populations of *P. australis* lies in their regenerative capabilities. In regeneration experiments, invasive populations demonstrated a significantly higher culm regeneration rate compared to source populations from Europe (*14*). In some cases, the invasive populations also exhibited greater rhizome regeneration, further enhancing their ability to colonize new areas and recover from environmental disturbances (*15*). This enhanced regenerative capacity likely provides invasive populations with a competitive edge, allowing them to establish dense monocultures that displace native species and transform ecosystem dynamics. Based on flow cytometry, North American populations of *P. australis* tend to have slightly larger genomes than their European counterparts (*16*) (*17*). While the precise reasons for the increased genome size remain unclear, emerging evidence suggests that the accumulation of B chromosomes, and a large number of tandem duplications may play a central role (*18*). B chromosomes are supernumerary chromosomes that exist alongside the standard set of A chromosomes, often without being essential for survival and regarded as parasitic(*19*). B chromosomes are known to carry large amounts of repetitive elements and non-coding ribosomal DNA (rDNA), and although they were traditionally thought to be genetically inert, recent studies suggest that they may actively participate in gene regulation, potentially causing deleterious effects (*20*). However, the functional role of B chromosomes in *P. australis* populations and the mechanisms by which they interact with the broader genome are still poorly understood.

In this study, we aimed to investigate how shoot systems regenerated from rhizomes contribute to the invasion success of *P. australis* in North America. To address this, we sampled the newly formed shoot tissues regenerated from rhizome and performed single-cell and spatial transcriptomic analyses for six samples maintained in a common garden. By analyzing gene expression patterns at both the cellular and tissue levels, we aimed to identify genes involved in the developmental process and compare their expression between invasive and source populations. Secondly, we aimed to investigate B chromosomecopy number variation and the evolution of invasion related genes using whole-genome and bulk RNA-seq data from the same individuals. Our goal was to uncover key genes involved in rhizome regeneration and their regulation by B chromosomes to better understand the genetic mechanisms underlying *P. australis* invasiveness.

## Results

### Single cell RNAseq data for common reed

The scRNAseq experiment yielded a comprehensive single-cell atlas for the shoot system of common reed (**Supplementary Fig. S1, Fig. 1a**). The number of captured cells across samples ranged from 7,154 to 15,297. The EU620 sample exhibited the highest median gene count, with 3,614 genes per cell, while the NAint61 sample had a median of 2,690 genes per cell (**Table 1, Supplementary Fig. S2**). After removing doublets and cells exhibiting high levels of mitochondrial and chloroplast expression and doublets, a total of 57,217 cells were retained for further analysis (**Supplementary Fig. S3**).

**Figure 1.**
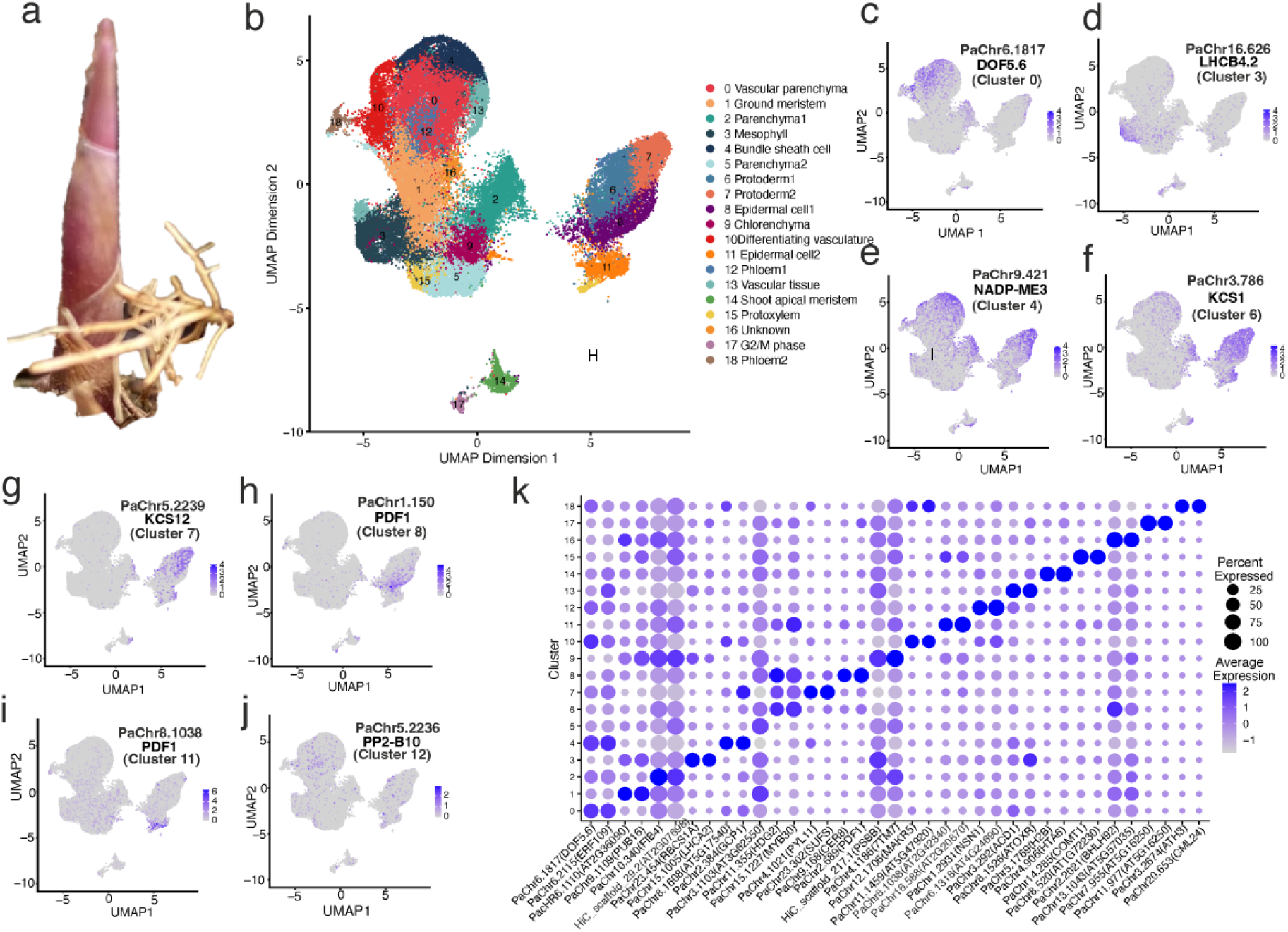
**a.** Tissue from common reed shoots was collected for single-cell and spatial transcriptomic analyses. **b.** Uniform Manifold Approximation and Projection (UMAP) plot showing dimensionality reduction of six single-cell samples following batch correction using the Harmony algorithm. The projection is based on 35 principal components (PCs) derived from highly variable genes. Nineteen distinct clusters are depicted in different colors. **c–j.** Expression levels of representative marker genes for Clusters 0, 3, 4, 6, 7, 8, 11, and 12. **k.** Bubble plot displaying the top two most reliable marker genes per cluster. Markers were identified using the FindAllMarkers function in Seurat, with the ROC test used to evaluate classification power. Markers were ranked by descending ROC score, followed by average log2 fold-change (log_2_FC) and cluster-specific expression frequency. Darker blue indicates higher log_2_FC values, while bubble size corresponds to relative expression levels within each cluster.

**Table 1.**
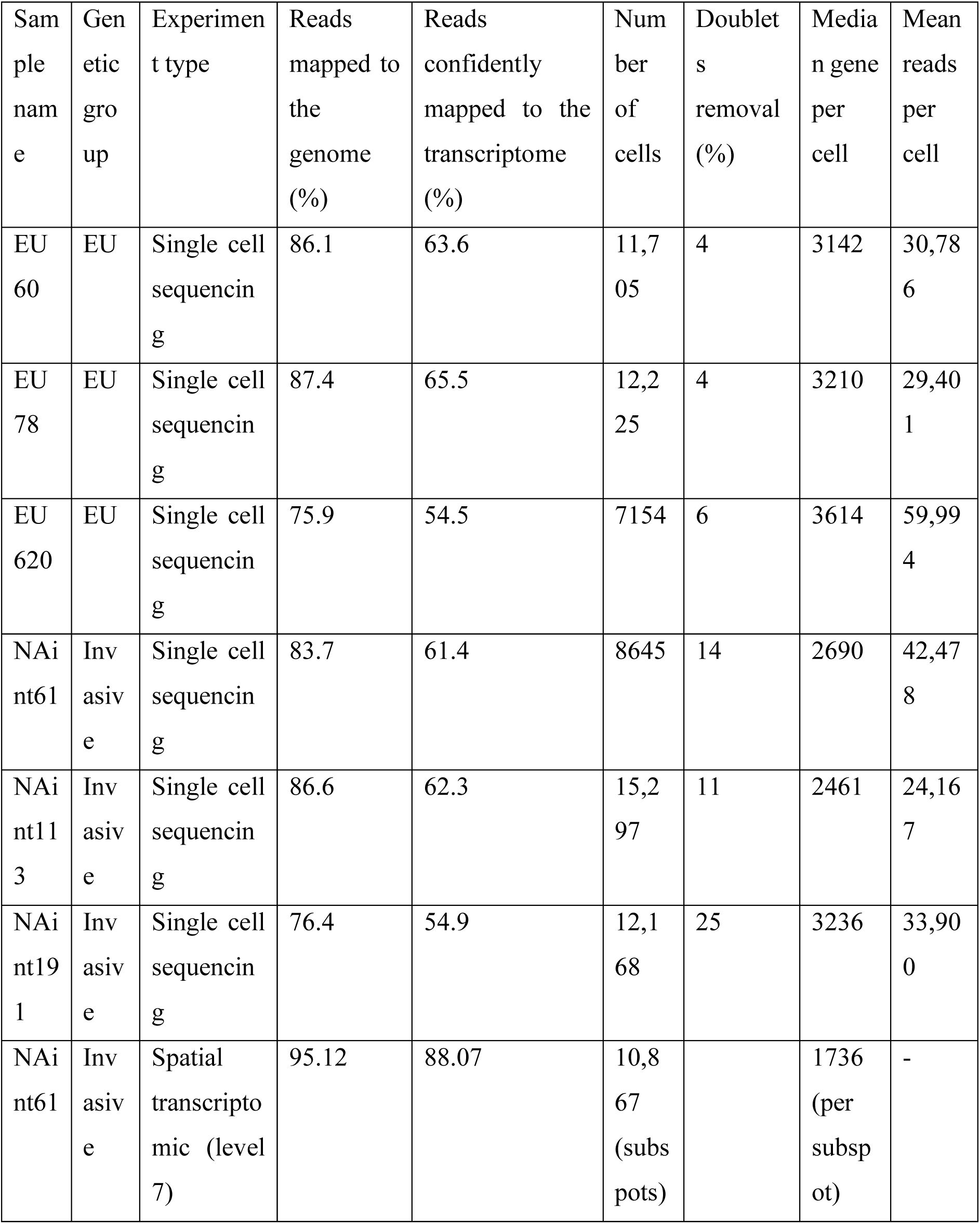
Sample information and sequencing quality for the single cell transcriptomic dataset.

### Single cell clustering

Using a resolution of 0.8, we identified 19 transcriptionally distinct cell clusters when visualized through Uniform Manifold Approximation and Projection (UMAP) and t-distributed Stochastic Neighbor Embedding (t-SNE) plots (**Fig. 1b, Supplementary Fig. S4**). Cluster-specific marker genes can be visualized on the UMAP (**Fig. 1c–j**) and a heatmap (**Fig. 1k**), and listed in **Supplementary Table S1.** Cell counts per cluster across the six *P. australis* samples from European (EU) and North American invasive (NAint) populations are summarized in **Supplementary Table S2**. The visualization of cell distribution for each sample on the UMAP can be found in **Supplemental Fig. S5**. Overall, Cluster 0 consistently exhibited the highest number of cells in both EU and NAint samples. Upon aggregating data from all six samples, we observed that Clusters 2, 5, and 9 had the lowest number of genes expressed per cell, indicating relatively low transcriptional activity. In contrast, Clusters 14 and 17 displayed the highest gene expression levels per cell, suggesting heightened cellular activity (**Fig. 2a**).

**Figure 2.**
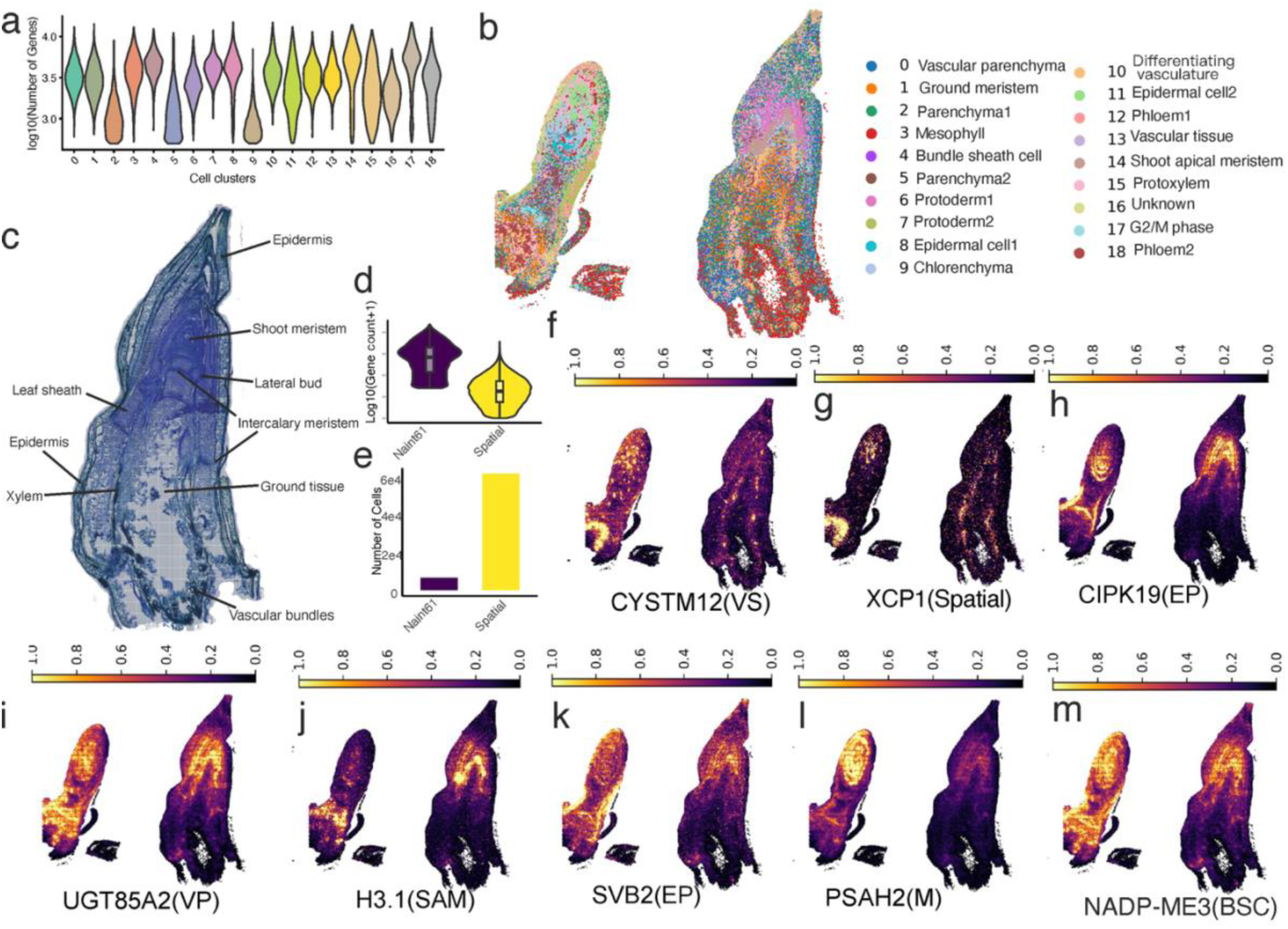
**a.** Violin plot showing the distribution of the number of detected genes (log₁₀-transformed) across individual cell clusters. **b.** Mapping of single-cell clusters (from six samples) onto spatial transcriptomic data (sample NAint61) using the Tangram algorithm. The color legend indicates the corresponding single-cell clusters, with spatial localization reflecting tissue-type associations. **c.** Anatomical structure of the shoot from spatial sample NAint61, including epidermis, shoot apical meristem, lateral bud, intercalary meristem, leaf sheath, xylem, vascular bundles, and undifferentiated ground tissue. **d.** Comparison of gene detection distributions between single-cell RNA sequencing and spatial transcriptomics (matrix generated using the cell split method) in sample NAint61. **e.**Comparison of cell number distributions between single-cell RNA sequencing and spatial transcriptomics in sample NAint61. **f–m.** Spatial expression patterns of marker genes corresponding to single-cell clusters, projected onto the spatial transcriptomic section. **Abbreviations: VS**, vascular tissue; **EP**, epidermal cells; **VP**, vascular parenchyma; **SAM**, shoot apical meristem; **M**, mesophyll; **BSC**, bundle sheath cells.

### Spatial transcriptome of a shoot tissue

Buds from the NAint61 individual were sectioned for spatial transcriptomics analysis (**Fig. 2b,c**). Anatomical sectioning of the large bud revealed eight distinct cell types along the vertical plane, including the epidermis, shoot apical meristem, lateral bud, intercalary meristem, ground tissue, vascular bundles, leaf sheath, and differentiating vasculature (**Fig. 2b,c**). The bud on the left side of the section offered a different perspective of the shoot, with its apical region positioned closer to the transection, enabling the identification of young leaf primordia and associated vascular tissues. We conducted a multiresolution analysis by aggregating different numbers of spatial barcoding spots into nine levels of subspots, each serving as an analytical unit. Summary statistics including the number of subspots, median UMI counts per subspot, median gene counts per subspot, and the total number of detected genes,are presented in **Supplementary Table S3**. In comparison, the single-cell RNA-seq dataset yielded a median gene count per cell ranging from 2,461 to 3,614 (**Fig. 2a**). To integrate spatial resolution with cellular identity, we generated a cell-segmented dataset based on high-resolution imaging. This approach identified 78,448 cells, with a median of 332 genes per cell and a median UMI count of 372. Although the number of genes per cell is lower than in the single-cell dataset, this method enabled the capture of a substantially larger number of cells (**Fig. 2d,e**). The highest gene expression levels were observed in young, actively growing tissues, such as meristematic regions and protoderms (**Supplementary Fig. S6**). Spatial transcriptomics analysis at subspot level 7 identified 19 distinct spatial clusters (**Supplementary Fig. S7**). To map transcriptional profiles onto anatomical structures, gene expression signatures from the single-cell RNA-seq clusters were projected onto the spatial transcriptomic dataset (**Fig. 2b; Supplementary Fig. S8**). Spatial cluster-specific marker genes were identified and summarized in **Supplementary Table S4**.

### Cell type identification from single-cell data and its spatial projection

We annotated the cell types of each single-cell cluster based on molecular markers reported in the literature (**Table 2; Supplementary Table S2**). To further validate spatial identity, representative marker genes from each single-cell cluster were projected onto the spatial transcriptomic data, as shown in **Fig. 2f–m**.

**Table 2.** Classical markers for each cell type.

| Cluster | Cell type | Gene markers | Reference |
| --- | --- | --- | --- |
| 0 | Vascular parenchyma | DOF zinc finger protein (DOF5.6) | (21, 22) |
| 1 | Ground meristem |  |  |
| 2 | Parenchyma1 |  |  |
| 3 | Mesophyll | *ribulose biphosphate carboxylase small chain 1A (RBCS1A)*, *chlorophyll a-b binding protein CP29.2 (LHCB4.2)*, *chlorophyll a-b binding protein CP26 (LHCB5)*, *light-harvesting chlorophyll B-binding 2 (LHCB2)*, *light-harvesting chlorophyll B-binding protein 3 (LHCB3)*, *photosystem I chlorophyll A/B-binding protein 1 (LHCA2)*, *photosystem I reaction center subunit III (PSAF)*, and *photosystem I subunit E-2 (PSAE-2)* | (22, 23) |
| 4 | Bundle sheath cells | *NADP-dependent malic enzyme 3 (NADP-ME3)*, *scr-like 23 (SCL23)*, *phenylalanine ammonia-lyase1 (PAL1)* | (22) (24, 25) (26) (27) (28, 29) (30) |
| 5 | Parenchyma2 |  |  |
| 6 | Protoderm1 | *KCS1*, *KCS11*, *KCS12*, *FDH (KCS10)*, *homeodomain glabrous 2 (HDG2)* | (22, 31-33) (30) |
| 7 | Protoderm2 | *KCS1*, *KCS11*, *KCS12*, *FDH (KCS10)*, *AtLTP1*, *LACS4* | (22, 31-33) (34) (35) (23) |
| 8 | Epidermal cell1 | *KCS1*, *KCS11*, *KCS12*, *FDH (KCS10)*, *Arabidopsis protodermal factor 1 (PDF1)*, *hydroxysteroid dehydrogenase 1 (HSD1)* | (22, 31-33) (23, 30, 36) |
| 9 | Chlorenchyma |  |  |
| 10 | Differentiating vasculature | *ATHB-5* | (23, 37) |
| 11 | Epidermal cell2 | *KCS1, KCS11, KCS12, FDH (KCS10), Arabidopsis protodermal factor 1 (PDF1), long-chain acyl-CoA synthetase 2 (LACS2)* | (22, 31-33)<br>(23, 30, 36) |
| 12 | Phloem1 | *phloem-specific marker PHLOEM PROTEIN 2-B10 (PP2-B10)* | (38) |
| 13 | Vascular tissue | *ATHB-7* | (23, 37) |
| 14 | Shoot apical meristem | *high mobility group B protein 1 (AtHMGB1)* | (39) |
| 15 | Protoxylem |  |  |
| 16 | Unknown |  |  |
| 17 | G2/M phase | *cyclin-dependent kinase (CDKB)* | (40) |
| 18 | Phloem | *DOF5.3* | (41) |

To gain further insight into cluster identity, we performed Gene Ontology (GO) enrichment analysis based on cell markers and analyzed the spatial locations for each single-cell cluster (**Supplementary Fig. S9**). For Cluster 1, enriched GO terms were related to responses to oxygen-containing compounds, abiotic stimuli, lipids, and hormones. Spatial projection placed these cells in the ground tissues surrounding intercalary meristems, suggesting they are ground mesristems. Cluster 2 and 5, located beneath Cluster 1 in leaf sheaths, ground tissue, and epidermis in the spatial projection, with only one to five markers discovered, likely represents parenchyma tissues. Cluster 9 had few markers but projected to mesophyll regions, and trajectory analysis supported its identity as developing chlorenchyma. Cluster 10 was mapped to differentiating vascular tissue in the spatial data and also exhibited expression of vascular marker genes. The gene markers of Cluster 14 are enriched in biological processes such as translation, negative regulation of DNA recombination, nucleosome assembly, macromolecule biosynthesis, negative regulation of DNA metabolic processes, protein-DNA complex assembly, chromatin organization, heterochromatin organization, organonitrogen compound biosynthesis, cell division, cellular biosynthetic processes, DNA dealkylation, DNA demethylation, and RNA metabolism, suggesting an active role in cell proliferation. Its spatial projection to the shoot meristem supported its annotation as the shoot apical meristem (SAM). Cluster 15 exhibited GO enrichment in the S-adenosylmethionine metabolic process and was projected to the vascular system. Given the known role of S-adenosylmethionine in lignin biosynthesis, which is characteristic of lignified tissues, we classify this cluster as xylem (*42*).

Cluster 16, enriched in genes related to responses to external biotic stimuli, remains unclassified. Cluster 17 showed enrichment in GO terms related to translation, organonitrogen compound biosynthetic processes, cytoplasmic translation, ribonucleoprotein complex biogenesis, cellular component organization or biogenesis, regulation of the cell cycle, mitotic phase transition, translational elongation, cellular component assembly, regulation of DNA-templated DNA replication, and cytoskeleton organization. These functions suggest that Cluster 17 is a cell cycle active population with high proliferative potential. Besides, its spatial projection to the lateral bud meristem indicates a subgroup of this cluster may represent lateral bud meristem cells. Cluster 18, which was projected to vascular tissues and exhibited high similarity to companion cells in *Arabidopsis*, is confirmed as phloem cells.

### Developmental pathways

RNA velocity analysis, derived from the ratio of spliced to unspliced transcripts, was used to infer dynamic cell state transitions. Using the apical meristem (Cluster 14) as the starting cluster, Clusters 18 and 11 consistently emerged as terminal states in 50% and 33% of the samples in the cell lineage trajectory respectively. Given that developmental programs are typically conserved within a species, we selected EU620 as a representative sample to explore cell type specific gene drivers. Cell fate mapping using CellRank identified three clusters with high fate probabilities (**Supplementary Fig. S10**), prominently including Clusters 1, 11, and three subpopulations within Cluster 9. These clusters correspond to ground meristem, epidermal cells, and parenchyma tissues, respectively. The identification of three distinct subgroups within Cluster 9 suggests either a degree of cellular heterogeneity or multipotent differentiation potential within this cluster.

The top key driver genes in the development of cluster 11 includes *PDF1 (Protodermal factor 1)*, *LTP3 (Lipid transfer protein 3)*, and *CASPL2A1 (Casp-like protein 2A1)* which peaked their expression at the onset of the differentiation, and *AT2G05540* (Glycine-rich protein family) which peaked its expression at the turning point (**Fig. 3a**). Pseudotime analysis of Cluster 11 revealed a sequential upregulation of lipid biosynthesis regulators (*MYB94*, *FAR4*, *KCS2*) to produce cuticular wax, initiators of trichome branching (*CML42*), adjustment of lipid compositions in the epidermis (*SSI2*, *ABCG11*, *LTPG1*), metal ion transportation (*CER-26-LIKE*, *ATHMP35*), and activation of vascular system fate genes (*PRX72*) as well as plant stuctural development regulators (*AOC3*) (**Fig. 3b**). The expression of the sequential genes also progressively expressed from multiple cell clusters to more localized in Cluster 11 (**Fig. 3e**). A few paralogues genes from different subgenomes functions synergistically in different time points, indicating a subfunctionization or neofunctionization during the rediploidization. In contrast, the top drivers of Cluster 9 included *MT2B*, *ATHCYSTM12*, *NRX1*, *AT3G17020* at early stage (cell state 1), *FIB4*, *DGK1*, *nPAP* and *LRK10L-2.8* in the intermediate stage (cell state 3), and *PSBB*, *RBCL*, *PSBC* and *LHB1B2* at the late stage (cell state 2) in the pseudotime estimation (**Supplementary Fig. S11**). Since the cell state 2 in cluster 9 showed high probability with determining the cell type, we focus on this subcluster there.

**Figure 3.**
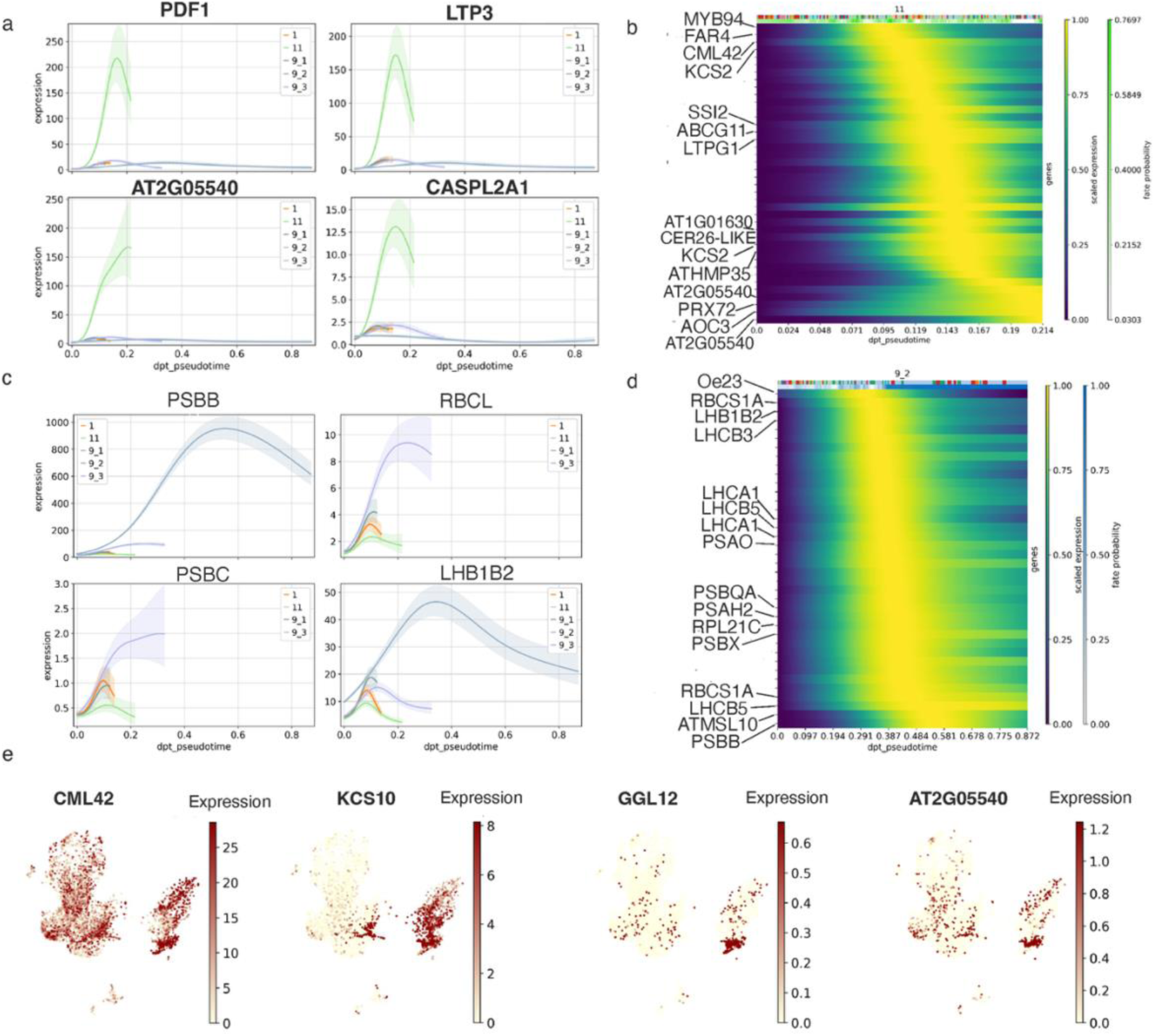
**a.** Dynamic expression patterns of key driver genes in Cluster 11 across its developmental trajectory, including *PaChr8.1038 (PDF1)*, *Chr10.1228 (LTP3)*, *PaChr6.1298 (AT2G05540)*, and *PaChr11.1614 (CASPL2A1)*. **b.** Representative gene expression profiles along the pseudotime trajectory, highlighting cell fate determination genes using Cluster 11 as a terminal state. The yellow shaded region indicates the time window of peak gene expression. **c.** Top driver genes in Cluster 9 exhibit peak expression at distinct stages along its pseudotime developmental trajectory. **d.** Temporal expression dynamics of genes associated with cell fate determination in Cluster 9 (cell state 2) along the pseudotime trajectory. The yellow highlight indicates the point of peak expression, while dark blue shading represents the probability of the gene’s involvement in fate determination. **e.** UMAP plots showing spatial expression of sequentially activated cell determination genes during the development of Cluster 11: *PaChr3.1219 (CML42)*, *PaChr3.474 (KCS10)*, *PaChr13.837 (GGL12)*, and *PaChr6.1299 (AT2G05540)*.

In Cluster 9, cell state 2 retained expression of numerous genes encoding photosystem components, which are well-established mesophyll markers. These genes also exhibited high cell fate probabilities (**Fig. 3c,d**), suggesting a strong commitment to mesophyll identity. Given the continuous developmental trajectory observed within Cluster 9 toward mesophyll fate, we propose that Cluster 9 represents the chlorenchyma tissue in *P. australis*.

### Differentially expressed genes between EU and invasive lineage

When comparing single-cell transcriptomic data between European and North American invasive populations, we observed that the invasive populations generally exhibited a slightly lower median cell counts (**Supplementary Table S1**; **Supplementary Fig. S12**). To ensure robust differential expression analysis, we pooled the single-cell RNA counts of each cluster and created pseudobulk RNA counts per sample. This approach reduces the number of multiple comparisons across thousands of cells and thus minimizes false positives (*43*). While PCA analysis of the pseudobulk expression profiles could distinguish invasive from non-invasive groups, a substantial proportion of variation was still observed within populations. This pattern shows a slight bias compared to the PCA results derived from WGS variant data and bulk RNA-seq of leaf tissue, where the largest proportion of variation was observed between groups (**Fig. 4a, b**; **Supplementary Fig. S13**), suggesting the presence of batch effects or biological heterogeneity within groups. Further supporting this interpretation, RNA velocity analysis indicated that sample EU620 exhibited transcriptional dynamics different from other European samples (**Supplementary Fig. S14**). The gene expression of non-invasive EU620 seems to be more correlated with another two invasive samples, suggesting potential technical noise such as high ratio of doublets in this sample (**Supplementary Fig. S15**).

**Figure 4.**
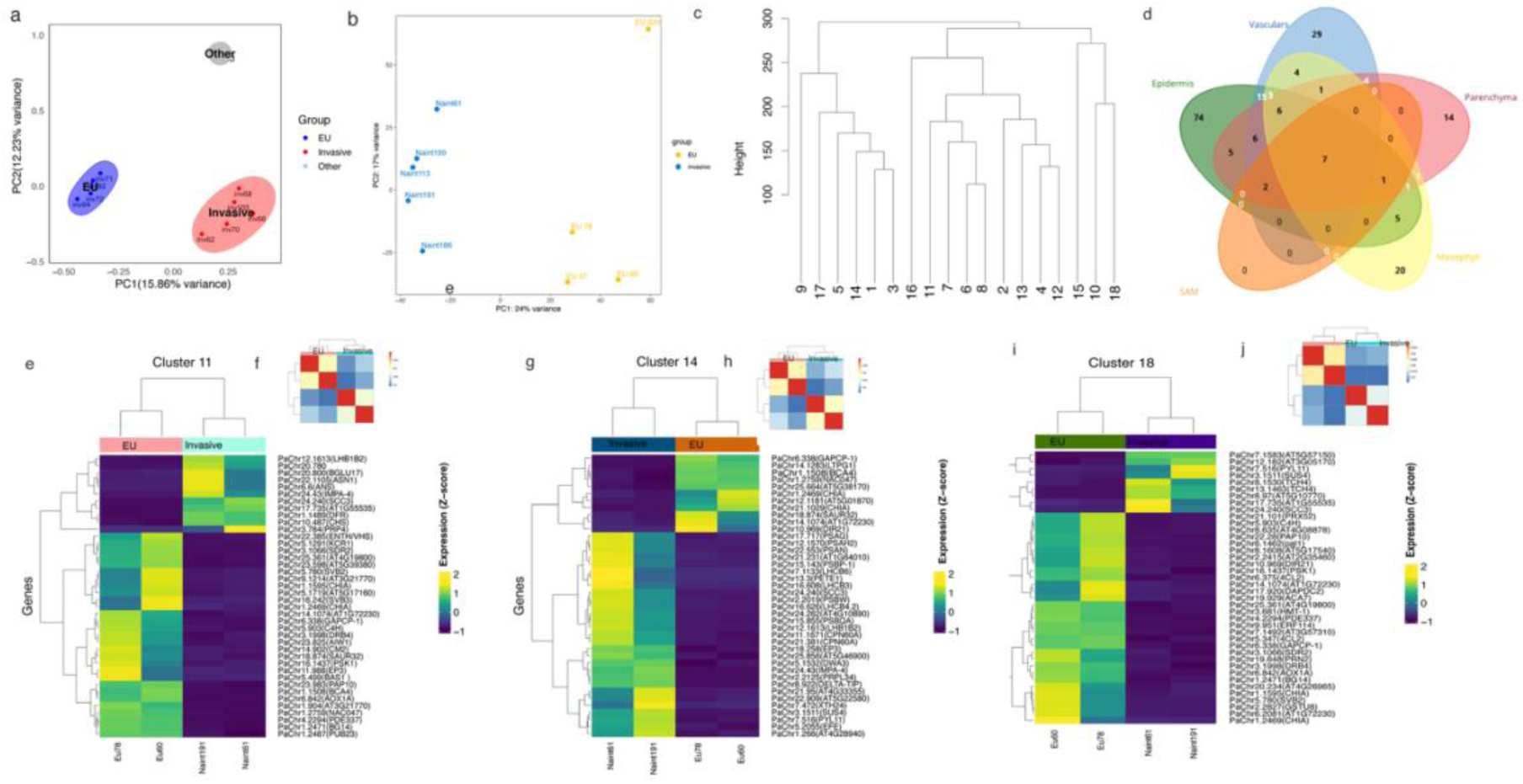
**a.** Principal Component Analysis (PCA) plot based on SNP variants obtained by aligning short reads from whole-genome sequencing to the reference genome. Samples enclosed in the blue ellipse represent European (EU) populations, while those in the red ellipse represent invasive populations. An additional sample, not enclosed, corresponds to a whole-genome sequence from an invasive individual reported in Oh et al. (2021). **b.** PCA plot based on RNA-seq read counts from leaf tissues mapped to the reference genome. A significant portion of the variance is explained by the separation between population groups, although gene expression variability is also observed within the EU group. **c.** Dendrogram depicting the relatedness of single-cell clusters, constructed using the hclust function with the complete linkage method. **d.** Venn diagram showing genes upregulated in the invasive population across five tissue types across six samples: vascular tissue, parenchyma, epidermis, shoot apical meristem (SAM), and mesophyll. Vascular tissues include Clusters 4, 10, 12, 13, 15, and 18; epidermal tissue includes Clusters 6, 7, 8, and 11; parenchyma includes Clusters 2, 5, 9, and 16; SAM corresponds to Cluster 14; mesophyll corresponds to Cluster 3. **f, h, j.** Heatmaps of the top 40 differentially expressed genes based on pseudobulk RNAseq between EU and invasive populations for Clusters 11, 14, and 18, respectively. **e, g, i.** Heatmaps showing sample-to-sample correlation based on pseudobulk RNA-seq data for the corresponding clusters.

Using correlations in gene expression among closely related cell types, we constructed a dendrogram to visualize their relationships based on a gene-by-cell-type matrix of average relative abundance (**Fig. 4c**). The topology may vary with different individuals, but cluster 6,7,8 and 11, as well as cluster 1 and 3 are stably grouped in the same clusters. We assigned each cluster to broader tissue categories for differential gene expression analysis: parenchyma (Clusters 2, 5, 9, 16), epidermis (Clusters 6, 7, 8, 11), vascular tissue (Clusters 4, 10, 12, 13, 15, 18), shoot apical meristem (Cluster 14) and mesophylls (Cluster 3).

Using three samples for each group, we observed the highest number of DEGs in the epidermis, with 37 genes upregulated, 16 of which (43%) are located on the B chromosome, and 89 genes downregulated in the invasive population (**Supplementary Tables S5, S6, S7**). The vascular system exhibited a slightly lower number of DEGs, with 21 genes upregulated, including six (29%) on the B chromosome, and 57 genes downregulated in the invasive samples compared to the European population. More DEGs were detected in undifferentiated than well-differentiated vascular tissues (**Supplementary Table S5**). In mesophyll cells from cluster 3, 19 genes were upregulated, nine of them (47%) on the B chromosomes, and 31 genes were downregulated in the North American introduced population (**Fig. 4d**). The commonly upregulated DEGs shared by the five tissues are *LRR4*, *SCC3*, *GATA2*, and a gene coding for *bHLH* superfamily protein. Apart from genes shared across multiple tissues, one gene encoding a Knotted1-binding protein, implicated in the establishment and maintenance of meristems in Poaceae species, was found to be upregulated in the invasive population in both epidermal and vascular tissues (**Supplementary Table S6**).

Concerning the high within group variability, we tentatively excluded outlier samples and retained the two most consistent replicates from each population for further hypothesis exploration. Among these, the vascular tissue exhibited the highest number of DEGs, with 749 genes upregulated and 728 genes downregulated in the invasive individuals. In the epidermis, 671 genes were upregulated and 697 were downregulated. The parenchyma displayed fewer DEGs, with 283 genes upregulated and 373 downregulated (**Supplementary Table S7**). Across all five tissue types, 45 genes were commonly upregulated in the invasive individuals. These genes were significantly enriched for GO terms related to photosynthetic light harvesting and the regulation of precursor metabolite and energy generation. In contrast, 27 genes were consistently downregulated across all tissues, but no significant GO term enrichment was identified for this group.

Excluding mesophyll cells which showed the fewest DEGs between groups, a total of 102 genes were commonly upregulated across the remaining four tissues, enriched in the light reaction of photosynthesis, including processes such as response to light stimulus, light harvesting in photosystem I and II, photosystem II assembly and stabilization, and the photosynthetic electron transport chain (**Supplementary Table S8**). Ninety-four genes were downregulated, with enrichment observed in respiratory burst involved in the defense response (**Supplementary Table S9**). The largest number of shared DEGs were detected between the epidermis and vascular systems, with 344 upregulated genes enriched in processes related to photosynthesis, stress responses (including hypoxia, low light intensity, heat, and cold), and brassinosteroid and glucosinolate metabolic processes (**Supplementary Table S10**). Specifically, the invasive population upregulated 36 genes in response to hypoxia, 31 genes in response to heat, 26 genes in response to cold, and 20 genes in response to light intensity. Additionally, 385 downregulated genes were enriched in immune system processes, including defense responses to bacteria and fungi, respiratory burst in defense, response to wounding, and various hormonal responses (abscisic acid, salicylic acid), as well as processes related to leaf senescence (**Supplementary Table S11**).

The upregulated genes exclusively in epidermis was enriched in processes related to response to brassinosteroids, fatty acids, jasmonic acid, oxidative stress, and endogenous stimuli. Biosynthetic processes such as flavonoid biosynthesis and monocarboxylic acid biosynthesis, as well as metabolic processes like xyloglucan metabolism, carboxylic acid metabolism, and olefinic compound metabolism, were also uniquely enriched in the epidermis. In contrast, the vascular tissue exhibited specific enrichment in response to mannose, response to hydrogen peroxide, cellular response to sulfur starvation, and cellular response to chemical stimulus (**Supplementary Table S12; Supplementary Note 1**).

Since Cluster 11, Cluster 14, and Cluster 18 exhibited the most prominent gene expression differences between the invasive and non-invasive groups, we selected the top 40 genes with the lowest adjusted p-values from these clusters for visualization (**Fig. 4e-h; Supplementary Note 2**).

### DEG analysis for the mature leaves between the two groups

BulkRNAseq data obtained from the mature leaf tissue of the same individuals clearly separated the European and North American introduced population in the first axis of PCA (**Fig. 4b**), although the EU620 again was an outlier for the European population. We found 147 genes were significantly upregulated in the invasive lineage, among which 68 (46%) were from the B chromosome (**Supplementary Table S13, S14**), and four genes were found to enrich in the biological process “fusion of sperm to egg plasma membrane involved in double fertilization forming a zygote and endosperm”. Ninety-seven genes were downregulated in the invasive population, with significant enrichment in multiple biosynthetic pathways, including isoprenoid, hydrocarbon, terpenoid, sesquiterpenoid, and diterpenoid biosynthesis, as well as biological processes such as response to herbivores and defense response to other organisms.

### B chromosome copy number variation and its influence on gene expression profiles

After mapping RAD-seq reads from 88 global individuals to the reference genome, we found that invasive individuals consistently exhibited a higher read depth ratio of B chromosomes to autosomes compared to other populations (**Fig. 5a**). This observation was further confirmed through whole-genome sequencing f nine individuals, including those used for single-cell and bulk transcriptomic analyses. The B chromosome-to-autosome read depth ratio averaged 1.99 ± 0.335 in the invasive lineages, indicating an average of four B chromosome copies (**Supplementary Table S15**). In contrast, the EU population exhibited a ratio of 0.261 ± 0.0115, suggesting a near absence of B chromosomes (**Fig. 5b**). Sliding window-based analysis of *Fst* comparison revealed exceptionally high divergence across the B chromosome, indicating strong genetic differentiation between the two populations (**Supplementary Fig. S16; Fig. 5c**). Tajima’s D revealed that the invasive population exhibited significantly lower (i.e., more negative) values compared to the EU population across all chromosomes (**Supplementary Fig. S17**), including the B chromosome (**Fig. 5d**), suggesting the influence of positive selection.

**Figure 5.**
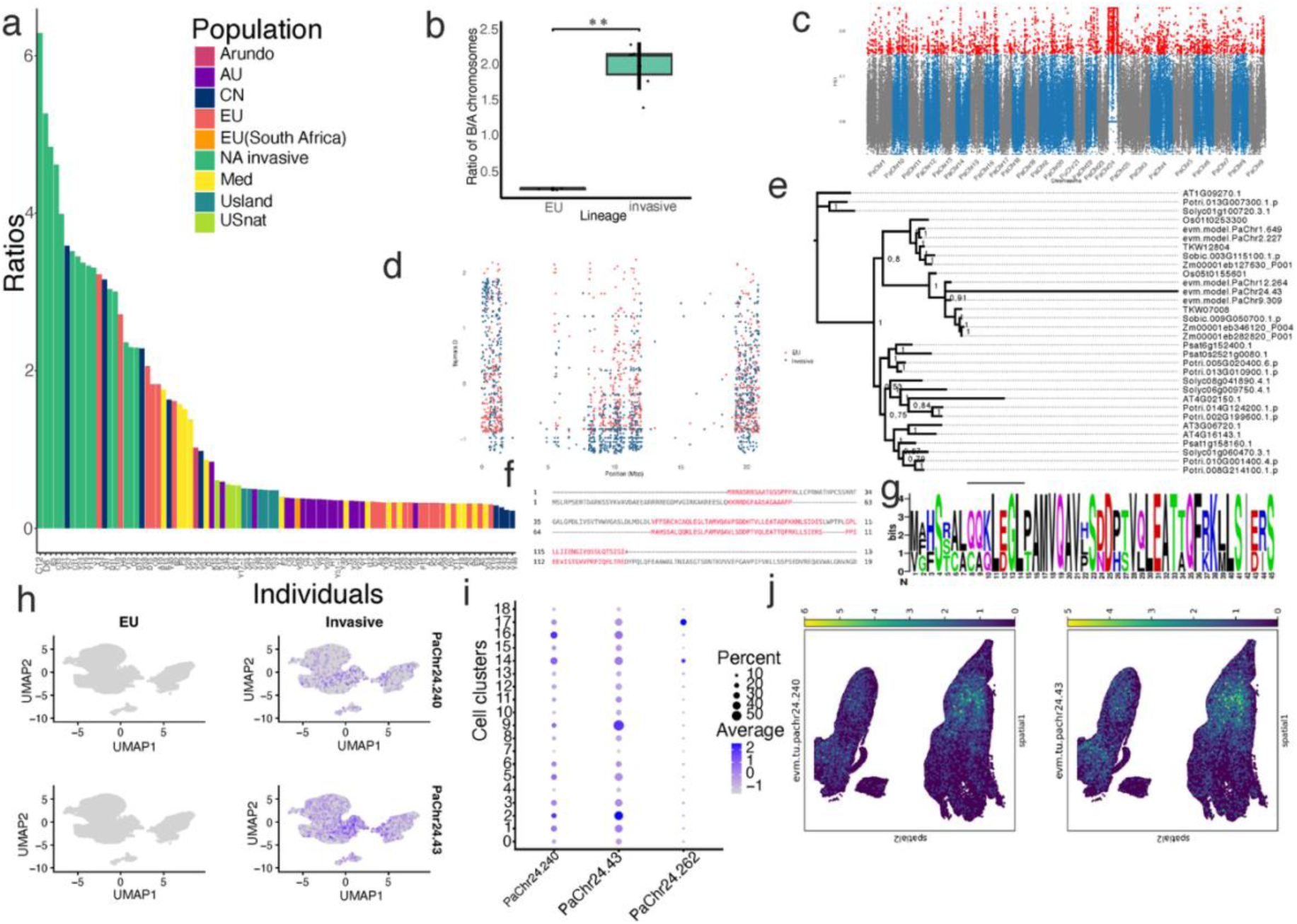
**a.** Bar plot showing the copy number of B chromosomes across different genetic lineages of *Phragmites australis*. Dark green bars represent invasive individuals in North America, all of which exhibit higher B chromosome copy numbers compared to other groups. **b.** Comparison of the ratio of reads mapped to B chromosomes versus those mapped to A chromosomes between invasive and non-invasive European populations. Statistical significance was assessed using the Mann–Whitney U test. **c.** Manhattan plot of *Fst* values between the EU and invasive populations across each chromosome, using sliding windows with a window size of 10 kb and a step size of 5 kb. **d.** Scatterplot of Tajima’s D values for two populations across the B chromosome. Red dots represent the values for the EU population, and blue dots represent the values for the invasive population. **e.** Phylogenetic tree of orthologs of the B chromosome gene *PaChr24.43* (a homolog of *IMPA-3*) across 10 species, revealing a relatively high mutation rate in this gene compared to others. **f.** Pairwise alignment of homologous genes *PaChr24.43* (B chromosome) and *PaChr9.309* (A chromosome). **g.** Conserved amino acid regions of *evm.model.PaChr24.43* among homologous genes in *P. australis* and *Zea mays*. **h.** Expression levels of two B chromosome genes (*PaChr24.43* and *PaChr24.240*) in non-invasive versus invasive individuals. **i.** Bubble plot showing the average expression levels (log₂ fold-change) of three B chromosome genes across clusters. Darker blue indicates higher average expression, while bubble size reflects the proportion of cells in which the gene is expressed. **j.** Spatial projection of the B chromosome genes *PaChr24.240* and *PaChr24.43* onto tissue sections.

Through pseudobulk differential expression analysis across each cluster, we identified 28 out of 286 genes on the B chromosome that are upregulated in the invasive population and function in different cell types (**Supplementary Table S16**). Of these, nine genes overlapped with regions of high positive Tajima’s D values (>2), while fifteen genes overlapped with regions exhibiting low Tajima’s D values (<0.9). For example, genes located on the B chromosome, *PaChr24.43* (homologous to *IMPA-3*) and *PaChr24.240* showed significantly higher expression in the invasive lineage across all clusters. The gene *OMTF3*, which encodes a methyltransferase, and *ABI5*, a transcription factor involved in ABA signaling, were exclusively upregulated in the epidermal cells of Cluster 8. The gene *MER3*, encoding a DNA helicase, was specifically upregulated in the shoot apical meristem (Cluster 14). Genes such as *APP1*, *AT5G62550*, *SKI3*, *RPA70B*, and *CDPK6* were uniquely upregulated in Cluster 2. In contrast, a few genes, including *SCC3* (*PaChr24.240* and *PaChr24.218*), *IMPA-3* (*PaChr24.43*), and *GATA2* (*PaChr24.82*), were upregulated in almost all clusters, suggesting that these genes are widely utilized throughout the plant.

To investigate the evolution of B chromosome genes, we examined the orthologues from a phylogenetic tree that included other Poaceae species. The gene *PaChr24.43* clusters with genes from chromosomes 9 (*PaChr9.309*) and 12 (*PaChr12.264*), forming a cluster with a group of importins, including *AT4G02150* in *Arabidopsis*, *Os05t0155601* in rice, *Zm00001eb346120* in maize, and *TKW07008* in *Setaria viridis* (**Fig. 5e**). The long branch length indicates a fast evolutionary rate of *PaChr24.43*(**Fig. 5f-g**). *PaChr24.240* and *PaChr24.218*, both encoding *SCC3* gene involving kinetochores functions, are clustered together with *PaChr10.167* and *PaChr21.972*. *PaChr24.262*, encoding for DDE family endonuclease, is clustered with *PaChr17.331*. Phylogenetic evolution of another two B chromosome genes (*PaChr24.218* And *PaChr24.240*) can be found in **Supplementary Fig. S18**. Three genes located on the B chromosomes were expressed across all clusters when comparing the EU and invasive lineages. mostly in young tissues (**Fig. 5h-j**).

## Discussion

Similar to rice, the aerial shoot system and rhizome elongation in common reed are initiated from axillary buds, and were differentiated rapidly after initiaion by controling the angle of the buds (*44*). In this study we focus on the aerial shoot, and the anatomical structure confirmed the shoot development following the general trend of monocot species. The young leaves emerge from the shoot apical meristems, and leaf sheath start to emerge from the fourth or fifth leave (*45*). By integrating multiomics data, cell types including vascular tissues, parenchyma, shoot apical meristem, mesophyll, ground meristem, bundle sheath cell, phloem, protoxylem, protoderm, and epidermal cell were identified in the shoot sytem. Cell type specific markers aligned well with the known literatures on plant development, indicating the cell type specific genes are well conserved in embryophyte.

Development trajectories of several tissues infered from the scRNAseq dataset provided insights about the molecular regulation of tissue differentiation on chlorenchyma and epidermis. Chlorenchyma (Cluster 9) is a type of parenchyma tissue in plants that contains abundant chloroplasts, typically found in mesophyll of leaves, and plays a crucial role in photosynthesis by capturing light energy and synthesizing organic compounds. The trajectory prediction infered the cell states of chlorenchyma cells from early to more differentiated, covering a wide range of pseudotime, suggesting this cluster may possess totipotency, enabling them to differentiate into an entire plant (*46*). These findings align with a study on tomato callus using spatial transcriptomics, which showed that chlorenchyma cells contribute to a complete shoot system development in tomatoes (*47*). Apart from chlorenchyma, bundle sheath cells and vascular parenchyma being considered key sources of regeneration in Poaceae species(*48*). In contrast, the development of epidermal cell underwent typical developmental transitions from protoderms (Cluster 6 and 7) to more specific differentiated tissues (Cluster 8 and 11). Following the typical epidermal cell evelopment in plants, the cell fate determination and morphogenesis of epidermal cells was first regulated by a critical regulator *Protodermal Factor 2 (PDF2)* as seen in the cell markers of the protoderm clusters(*49*). Developmental trajectory of the mature epidermal cell (Cluster 11) showed the gene expression regarding to trichome branching occurs early, conforms with the notion that trichome differentiation may occur earlier than other stomata cells(*49*). More developmental process regarding aerenchyma formation are inferred from the spatial transcriptomics data. Spatial mapping of single-cell Cluster 1 to spatial clusters 4, 16, and 18 indicates that, in addition to its role as ground meristem, it is also likely associated with aerenchyma formation. In the marsh aquatic plant *Typha angustifolia* L. (Typhaceae), the development of leaf aerenchyma is divided into four stages based on morphological characteristics: solid, early cavity formation, late cavity formation, and mature cavity (*50*). Our data identified *APX1* as a cell-type marker for spatial Clusters 4. *Prx52*, a protein similar to peroxidases involved in lignin biosynthesis, was found to have three paralogues as markers for spatial clusters 16. These findings suggest that Cluster 1 in the single-cell data may initiate aerenchyma formation during the solid stage, as this stage is characterized by high expression levels of *APX* and *Prx* genes, which decrease as development progresses (*50*). The anatomical structure also revealed a looser tissue arrangement on the lower side of the shoot, consistent with the pith cavity formation process observed in common reed (*51*).

Developmental system drift, defined as the genetic divergence in the developmental pathways of homologous phenotypic traits over evolutionary trajectories(*52, 53*), was observed in shoot development between the two populations of common reed. This was reflected in differential gene expression across distinct cell types between groups, with a notable involvement of B chromosome genes in the invasive population. Such divergence in regulatory programs may underlie the observed developmental system drift between invasive and native populations. It’s worth noting that for the individual EU620 was grouped within EU population in the genomic analysis but displayed as an outlier in both single cell and bulk RNA expression, suggesting changes in gene regulatory network, mostly by epigenetic effect. Invasive populations of *P. australis* showed a tendancy of exhibiting elevated expression of genes associated with photosynthetic activity and enhanced tolerance to hypoxic conditions, while genes linked to acute defense responses are downregulated compared to non-invasive European populations. This pattern aligns with life history trade-off theory, which suggests that energetically costly defensive traits are often lost in favor of traits that support rapid growth and high fecundity. A similar shift has been observed in *Ageratina adenophora*, where invasive populations allocate more nitrogen to photosynthetic activities rather than structural cell construction, unlike their native counterparts (*54*).

Elevated photosynthetic rates have been reported in many invasive plant species relative to co-occurring native species, such as *Rubus* spp. (*55*), *Berberis darwinii* (*56*), and *Tithonia diversifolia* (*57*). Furthermore, invasive species such as *Mikania micrantha*, *Wedelia trilobata*, and *Ipomoea cairica* demonstrate more efficient CO₂ usage compared to native species(*58*). A comparative study of leaf traits across 92 pairs of co-distributed invasive and non-invasive species by Liu *et al.* concluded that invasive species typically evolve significantly higher leaf nitrogen concentrations, light-saturated photosynthetic rates, and efficiencies in the use of energy, nitrogen, phosphorus, and potassium for photosynthesis (*59*).

For common reed, invasive populations display significantly higher photosynthetic rates, stomatal conductance, and leaf nitrogen content than native North American populations, supporting the notion of enhanced photosynthetic capacity in the invasive lineage(*60*). In fact, the European source population already possessed more efficient photosynthetic machinery than the North American native population, suggesting a degree of preadaptation before the invasion(*61*). Moreover, the invasive population differs from its European ancestor in photosynthetic nitrogen use efficiency and structural investment costs (*61*). Despite these variations, the highest photosynthetic efficiencies are still found in populations originating from Mediterranean regions(*62*), implying that a broader array of genetic determinants, arising from long-term adaptation to tropical environments may further enhance photosynthetic performance.

Copy number variation of B chromosomes and their expression were found to be significantly associated with the bioinvasion process. Structural variations, such as large haplotypic block inversions, have been identified as key drivers of rapid adaptation to local environments in *Ambrosia artemisiifolia* (*63*). This haplotypic inversion preserves a region of adaptive loci in the invasive populations which includes selective sweep near the flowering locus *ELF3*, but not in native populations, providing a potential advantage for its invasion success. Similar structural variation mechanisms have also been observed in *Helianthus* species (*64*). Subsequently, structural variations may be transferred to other populations through hybridization, boosting their potential for survival and expansion (*65*). In our study, we observed variations in B chromosome copy number across invasive populations and other genetic lineages, indicating that B chromosomes may serve as hotspots for rapid adaptation. The more negative Tajima’s D values observed on the B chromosome are consistent with expectations, as its higher copy number in the invasive population likely contributes to an increased number of low-frequency SNPs.

The 28 upregulated genes on the B chromosome in the invasive populations are involved in several key functions, including DNA segregation and repair, gamete generation, methylation, photomorphogenesis, transposable ability, and plant hormone responses (such as auxin, brassinosteroids, and gibberellic acid), as well as responses to hypoxia. Ninety-three percent of these genes are expressed in both shoot and mature leaf tissues, suggesting the effect of B chromosome covers the whole life history of the plant. The genes largely overlap with regions showing either the highest or lowest Tajima’s D values, suggesting they are subject to both balancing selection and directional selection. The gene *PaChr24.43* is homologous to *IMPORTIN-α3/MOS6 (MODIFIER OF SNC1, 6)* in *Arabidopsis*, a key recruiter of nuclear localization signal-containing cargo proteins into the nucleus (*66*). *IMPA-3* is an importin involved in transporting essential proteins into the nucleus, while cohesin (*SCC3*) is believed to maintain the three-dimensional structure of plant genomes, particularly during mitosis and meiosis (39). The coordinated function of *IMPA-3* and *PaChr24.240* (homologous to LOC133886699, encoding a sister-chromatid cohesion protein 3-like protein) may help prevent chromosomal abnormalities during rapid cell division, thereby promoting enhanced cell proliferation, genome stability, and increased adaptability to environmental stress. *IMPA-3* functions as the main receptor for the R (resistance) gene *SNC1* in *Arabidopsis*, playing a critical role in plant innate immunity (*67, 68*). R genes are known to evolve rapidly, even within the same species (*69*). Interestingly, the importin protein on the B chromosome is also undergoing rapid evolution, as evidenced by the phylogenetic tree of this gene in *P. australis*. In contrast, its paralogue on Chromosome 9 does not exhibit rapid evolution but follows a typical evolutionary rate seen in other species. Additionally, the homologous gene *PaChr9.309* did not show differential expression between the two populations, whereas *PaChr24.43* showed significant expression differences across every cell type. This suggests that the *IMPA-3* gene on the B chromosome is under stronger selection and may be more highly favored in the invasive populations. Moreover, *PaChr24.43*, compared to the homologous *PaChr9.309* (which contains 531 amino acids), is truncated, with only 133 amino acids, indicating that it represents only a partial version of the gene, and has likely undergone neofunctionalization or subfunctionalization. This truncated form may have been translocated to the B chromosome through the activity of transposable elements. Coincidentally, *PaChr24.262*, homologous to *AT4G10890*, which encodes a DDE family endonuclease, was also a major upregulated gene in the invasive lineage. DDE superfamily genes are involved in coordinating metal ions for catalysis, enabling site-specific DNA cleavage and subsequent strand transfer (*70*), potentially working as a transposase protein that facilitates the formation of transposable elements in *P. australis* (*71*). Interestingly, DDE proteins have undergone neofunctionalization and are crucial regulators of heading date in rice, underscoring their versatile roles in plant development (*72*). *GATA2*, a transcriptional regulator involved in brassinosteroid and light signaling pathways, positively regulates photomorphogenesis in plants (*73*). The gene *PaChr24.82* is partially homologous to *Arabidopsis GATA2* (67 nucleotides) and encodes a protein sequence of 259 amino acids. Interestingly, we did not find orthologous genes from the plant species compared, suggesting that this gene may have evolved specifically on the B chromosome, acquiring a unique function.

Similar to other Poaceae species, B chromosome drive may also exist in *P. australis* to facilitate the non-Mendelian inheritance. The genes *PaChr24.218* and *PaChr24.240*, both homologous to the *Arabidopsis SCC3* gene, which is essential for the monopolar orientation of kinetochores during meiosis, are upregulated in the invasive population across nearly all cell types. Previous studies have linked genes encoding microtubule-associated proteins, which influence cell division to drive mechanism (*74*). The upregulation of *SCC3-like* genes in the invasive population may thus suggest the presence of a B chromosome drive mechanism. Additionally, the co-occurrence of duplicated homologous genes could indicate a synergistic effect or dosage compensation in the regulation of meiosis, further supporting the stability and persistence of B chromosomes in the invasive lineage.

## Materials and Method

### Tissue dissociation and preparation of single-cell suspensions

To generate a comprehensive single-cell and spatial transcriptomic atlas for *P. australis*, we selected six individuals from EU (non-invasive) and NAint (invasive) populations, with three biological replicates in each group. These plants were grown in the common garden of Shandong University under natural conditions. Prior to the single cell protoplast preparation, the plants were kept at room temperature (25 ℃) for a few days, and then the rhizome shoots were collected. From each individual, we collected between 5 and 22 buds, which were then pooled into a petri dish for each sample. The rhizome shoots were cut into 1-2 mm strips and added to tubes containing 10ml enzyme solution consisting of 2% cellulase R10, 0.8% macerozyme R-10, 0.5% pectinase Y-23, 0.1% BSA, 20mM MES,20mM KCl,10mMCaCL2 and 0.55M mannitol with shaking at 60rpm for 1 hour at 28℃. Cells were then filtered with a 40 um cell strainer. Cell activity was detected by trypan blue staining and cell concentration was measured using a hemocytometer and a light microscope. Lastly, protoplasts were resuspended in 8% (w/v) mannitol solution in preparation for loading onto the chromium controller of the 10xGenomics platform. Approximately, 1.6×104 isolated single cells and enzyme gel-beads were packed into a single oil droplet for single-cell RNA-seq library construction.

### Chromium 10x Genomics library and sequencing

Cells were loaded onto the 10X Chromium Single Cell Platform (10X Genomics) at a concentration of 700-1200cells/ul (Single Cell 3′ library and Gel Bead Kit v.3) as described in the manufacturer’s protocol. Generation of gel beads in emulsion (GEMs), barcoding, GEM-RT clean-up, complementary DNA amplification and library construction were all performed as per the manufacturer’s protocol. Qubit was used for library quantification before pooling. The final library pool was sequenced on the Illumina Nova6000 instrument by Shanghai Personal bio (Shanghai, China) using 150bp paired-end reads.

### Spatial transcriptomics sample preparation

We selected NAint61 from the individuals that were used for single cell analysis and performed spatial transcriptomics. A number of fresh shoots are frozen using isopentane and liquid nitrogen bath. Then the frozen tissue samples are embedded in optical cutting temperature (OCT) and stored at −80°C. Frozen sectioning is performed using a cryostat, and the tissue sections are collected for RNA extraction and quality control. We selected the samples with a minimum RNA Integrity Number (RIN) value of ≥6 to ensure that the RNA is not degraded during the tissue freezing process. Tissue optimization is performed on the samples before the formal experiment to determine the optimal permeabilization time for spatial gene expression library construction. For the gene expression library preparation, tissue sections are mounted on gene expression slides, followed by methanol fixation, cell segmentation fluorescence hematoxylin and eosin (H&E) staining, cell segmentation fluorescence imaging, H&E staining, and bright-field imaging. Then, tissue permeabilization is performed. The mRNA released from the tissue sections on the gene expression slides is captured by specific probes on the spots and reverse transcribed into cDNA. The cDNA with spatial barcodes is collected from the carrier slides. After double-stranded synthesis, denaturation, PCR amplification, the initial cDNA is completed. Subsequently, enzymatic fragmentation, end repair with A-tailing, magnetic bead fragment selection, adapter ligation, magnetic bead purification, and sample index PCR are performed to construct standard next-generation sequencing libraries. After passing library quality control, the spatial gene expression library is sequenced using a next-generation sequencing platform with a Pair-end 150 sequencing strategy.

### Sequencing and data preprocessing of scRNA data

We first demultiplexed the raw reads of scRNA and removed the barcodes of six individuals (EU60, EU78, EU620, NAint61, NAint113, and NAint191). Then the clean reads were mapped to the chromosome level genome assembly using cellranger pipeline v8.0 (*75*). The reference genome sequence of *P. australis* was obtained from our previous work (NCBI accession: ASM4037322v1) with a slight modification(*18*). Since the genome assembly spans from telomere to telomere, we consider it complete and replaced the remaining contigs with complete chloroplast and mitochondrial genomes. Annotation of the modified assembly was performed using Liftoff, based on the original gene models(*76*). The mkref function with default settings in CellRanger was used to build the reference for this species, and count function was used to assess the number of unique molecular identifiers (UMIs) for each gene and barcode. The count matrices were then used for quality control using scanpy(*77*). Only genes that are expressed in at least 20 cells, and cells that have at least 500 genes expressed were kept in the dataset. Cells with mitochondrial genes higher than 10% and chloroplast genes higher than 30% may indicate dead cells thus were removed from the analysis. Doublets were removed using DoubleFinder(*78*). The data of six samples was then integrated into one Seurat object using R package Seurat(*79*), and normalized using SCTransform V2. The normalized gene expression matrix that was further analyzed using the Seurat package for dimensionality reduction via PCA analyses. To correct for batch effects across samples, we applied the Harmony integration method (*80*). In total, 35 PCA axis were selected for further analysis. After trying with different resolution, we selected 0.8 as the final resolution, clustered the cells via the Louvain algorithm and run UMAP with parameter min.dist = 0.1, n.neighbors = 50.

### Differential gene expression of scRNA data

The differential gene expression of scRNA data between invasive and non-invasive group was performed using DEseq2(*81*). The read counts of all clusters were first aggregated for each sample to construct pseudobulks, transforming a genes-by-cells matrix to a genes-by-replicates matrix using matrix multiplication. Then, the counts were normalized using rlog method in DEseq2, and visualized using PCA. Differential expression comparisons were performed on the two groups. Since the intragroup variations accounted for most of the variations, we removed the outliers of EU620 and NAint113, only keeping two samples per population as replicates. The differential expression were then perfromed for each cluster using DEseq2, with a Wald test of the negative binomial model coefficients (DESeq2-Wald) to compute the statistical significance. A shinked log2 fold change (LFC) based on the apeglm method(*82*) was employed to improve the gene rankings of significantly differentiated genes.

### Data analysis of spatial transcriptomics

For the spatial transcriptomic data, we used subspot level 7 which includes 127 spots with a resolution of 37µm as a unit to proceed with the downstream analyses. The sequencing data was aligned and quantified using the BMKGENE official software, BSTMatrix. Read 2 was aligned to the reference genome using the STAR software. Cell split was also performed using BSTMatrix, based on a high-resolution image obtained during tissue sectioning. The expression matrix was used as input in Seurat and normalized with SCTransform V2 for downstream analysis, including running PCA, and UMAP (Uniform Manifold Approximation and Projection) visualization. The clustering was done with a resolution of 0.5, and spatial cluster-specific marker genes were identified using Seurat’s FindMarkers function.

### Cell type annotation and gene markers of single cell clusters

The cell clustering and processing was performed using Seurat R package(*83*). In total, 19 clusters were obtained. For cell type annotation, we first clustered the single cell experiments and projected them to the spatial transcriptomic data to find the matching tissues. Then, we used FindAllMarkers function in Seurat with a threshold of logfc.threshold = 1, min.pct = 0.1, only.pos = TRUE to find all the cell markers for each cluster. We searched for the GO terms of these marker genes to conclude the function of each cluster. Thirdly, we searched for orthologous genes for the cell markers in closely related plant species, including maize, rice, sorghum, and compared cell markers with other literature. To validate the annotation accuracy of the cell types, we projected the cells and clusters of single cell dataset to the spatial transcriptomic slides using both Seurat and Tangram(*84*). Tangram was said to be one of the most accurate software tools for the projection so far, as evidence by a benchmark study(*85*). The denrogram which groups the relevant single cell clusters were constructed using MuSiC(*86*).

### RNA velocity and trajectory analysis

Following preprocessing steps, the Seurat object of EU620 was converted to h5ad format, and the RNA velocity was evaluated by a velocyte(*87*) run. The loom file containing splicing information was generated with default parameters and RNA velocity was conducted with a python package scvelo(*88*). The terminal stages and pseudotime of cells were inferred using cellrank(*89*), with the meristematic cell cluster 14 set as the starting cluster.

### Whole genome sequencing

To evaluate the B chromosome number in population level, we reused the RADseq dataset of 88 individuals obtained from the previous published work(*10*). The reads were mapped to the reference genome with repeats masked, and the ratio of reads mapped to the B chromosome and other chromosomes were calculated. In addition, whole genome sequences were obtained from 10 individuals representing 5 individuals from European ancestral and 5 individuals from North American invasive populations. The DNA were extracted using CTAB method, and were sequenced using BGI DNBseq platform. The reads of RADseq and WGS datasets were cleaned using trimmomatic and mapped to the reference genome using BWA-mem (*90*). The alignment was first removed PCR duplicates using GATK MarkDuplicates (*91*), and then input into Deepvariant (*92*) followed by GLnexus (*93*) to call the single nucleotide polymorphisms (SNPs). To detect the structural variations between the two lineages, we used Delly which predicts structural variations based on the split reads, and gridss to detect the insertions, deletions, inversions, and duplications between the two populations. We then merged the results using Survivor merge function. The number of reads mapped to the B chromosome and other 24 chromosomes was calculated using bamcoverage in deeptools(*94*). The differences in ratios between populations were tested using Mann-Whitney U test, with p < 0.05 considered statistically significant. Tajima’s D was calculated using PopGenome R package(*95*).

### RNAseq for the leaf tissues

We harvested mature leaves from nine individuals grown in a common garden at Shandong University in June 2023 and used them for transcriptomic analysis, enabling a direct comparison of gene expression profiles. The genetic background was inferred from the published phylogeographic pattern(*10*). The total RNA was extracted using the RNAprep Pure Plant Kit (Tiangen, Beijing, China) according to the instructions provided by the manufacturer. RNA concentration and purity was measured using NanoDrop 2000 (Thermo Fisher Scientific, Wilmington, DE). RNA integrity were assessed using the RNA Nano 6000 Assay Kit of the Agilent Bioanalyzer 2100 system (Agilent Technologies, CA, USA). A total amount of 1 μg RNA per sample was used as input material for the preparations of RNA samples. Sequencing libraries were generated using Hieff NGS Ultima Dual-mode mRNA Library Prep Kit for Illumina following manufacturer’s recommendations and index codes were added to attribute sequences to each sample. The libraries were sequenced on an Illumina NovaSeq X Plus platform to generate 150 bp paired-end reads. Quality of the reads were checked using fastqc (https://www.bioinformatics.babraham.ac.uk/projects/fastqc/). Adapters were clipped and low quality bases (phred score < 20) were removed from the reads in 4 bp sliding windows using Trimmomatic (*96*). Genome assembly and annotation of *P. australis* was obtained from Clean reads were mapped to the reference genome using STAR aligner (*97*). After sorting using Samtools (*98*), the mapped reads were processed using StringTie (*99*) to obtain the gene and transcript count matrix. Principal component analysis (PCA) was conducted on differentially expressed genes among the treatment groups using DEseq2 following rlog transformation of gene counts, which represent the number of reads mapped to the reference genome (*100*).

DESeq2 provides statistical routines for determining differential expression in digital gene expression data using a model based on the negative binomial distribution. The resulting P values were adjusted using the Benjamini and Hochberg’s approach for controlling the false discovery rate. Genes with an adjusted P-value < 0.01 & Fold Change≥2 found by DESeq2 were assigned as differentially expressed. Gene Ontology (GO) enrichment assessment analyses sets of genes in the context of GO categories to determine whether certain functional categories are overrepresented or enriched within the background whole genome gene sets. Here we tested enriched GO terms for the differentially expressed genes (DEGs) using goatools (*101*), using a criterion of P <0.05 with Bonferroni correction.

### Orthologues among species and transcription factor prediction

Eight outgroup species, including *Arabidopsis* (TAIR 11), rice IRGSP1.0 (https://rapdb.dna.affrc.go.jp/download/irgsp1.html)(102), *Populus trichocarpa* v4.1(Phytozome)(*30*), *sorghum v3.1.1* (Phytozome)(*103*), *Pisum sativum* (*23, 104*), *Solanum lycopersicum* ITAG4.0 (Phytozome)(*105*), maize B73 v5 and *Setaria_viridis* V2 reference genomes at https://plants.ensembl.org/ (*106*) were included to find out the orthologues between species using Orthofinder(*107*). This allows us to obtain genetic markers of each cell type and compare the gene expression between monocots and dicots in the evolutionary trajectories. Transcription factors of the proteins in common reed genome were predicted from PlantRegMap (*108*). The alignment between B chromosome sequences were performed using Cobalt(*109*) in NCBI website, and the visualization of conserved amino acid homologues were conductd using weblogo (*110*).

## Data availability

All sequences generated in this study have been store in NCBI database. The single cell transcriptome of six samples and the spatial transcriptomic data were deposited in GEO database with the number GSE295468. The whole genome sequencing and RNAseq data were deposited in the SRA database with a project number PRJNA1254975.

## Author contributions

C.W., LL. Liu, and WH. Guo conceptualized and supervised the research. LL. Liu and HJ. S prepared the materials for scRNA-seq and spatial transcriptomics. C.W. performed data analysis with contributions from J.O. and MX. Y. HJ. S and LL. Liu analyzed the phenotypic traits. YF. C participated in recognizing shoot anatomical structures. C.W. and J.S. jointly discovered the genomic structural variations. C.W. wrote the manuscript, with contribution from J.O., J.S., LL. Liu, MX. Y, and WH. Guo in the revisions.

## Acknowledgement

We thank Dr. Iivari Kleino for valuable discussions on single-cell data analysis. We are grateful to the staff at BMK for their comprehensive assistance in obtaining high-quality tissue section slides. We also thank Ms. Wenyi Sheng for her help in caring for the plant materials in the greenhouse. We thank Pawel Jan Roszak and Yrjö Helariutta for their assistance with the identification of vascular markers. This work was supported by the Natural Science Foundation of Shandong Province (No. ZR2021QC119 to C. W. and No. ZR2024QC197 to L.L. Liu), the National Natural Science Foundation of China (No. 32470388 to L.L. Liu and U22A20558 to W.H. Guo) and Fundamental Research Funds of Chinese Academy of Forestry (Grant No. CAFYBB2023ZA004 to L.L. Lin). C.W., J.O., and J.S. acknowledge funding from TreeBio - Research council of Finland Centre of Excellence in Tree Biology (decision 346139). We also acknowledge CSC Finland for their support of computational resources.

