## Supplemental Figures for "Single-cell and spatial transcriptomics uncover the role of B chromosomes in driving plant invasiveness"

### Supplementary materials

**Supplementary Figure S1** Several shoots generated from rhizome of the same sample that were used to create single cell suspension for the individual EU60.

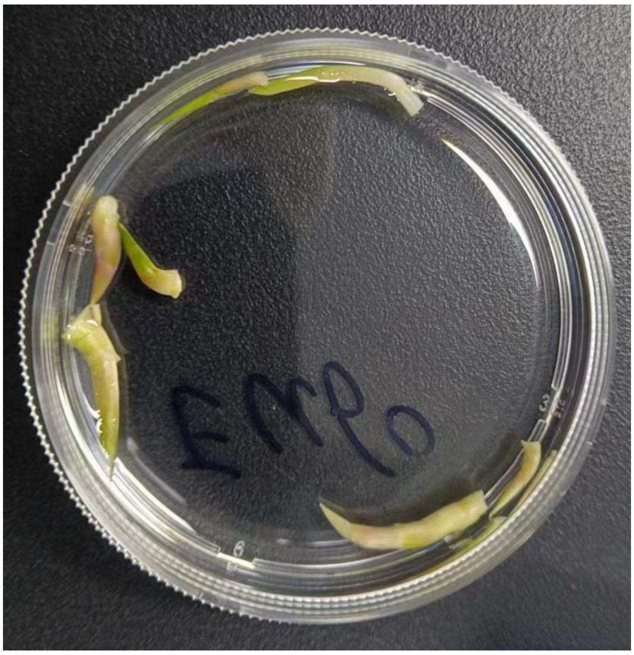

**Supplementary Figure S2 Violin plot showing the distribution of detected genes per sample.** Orange dots denote cells below the expression threshold (excluded during QC).

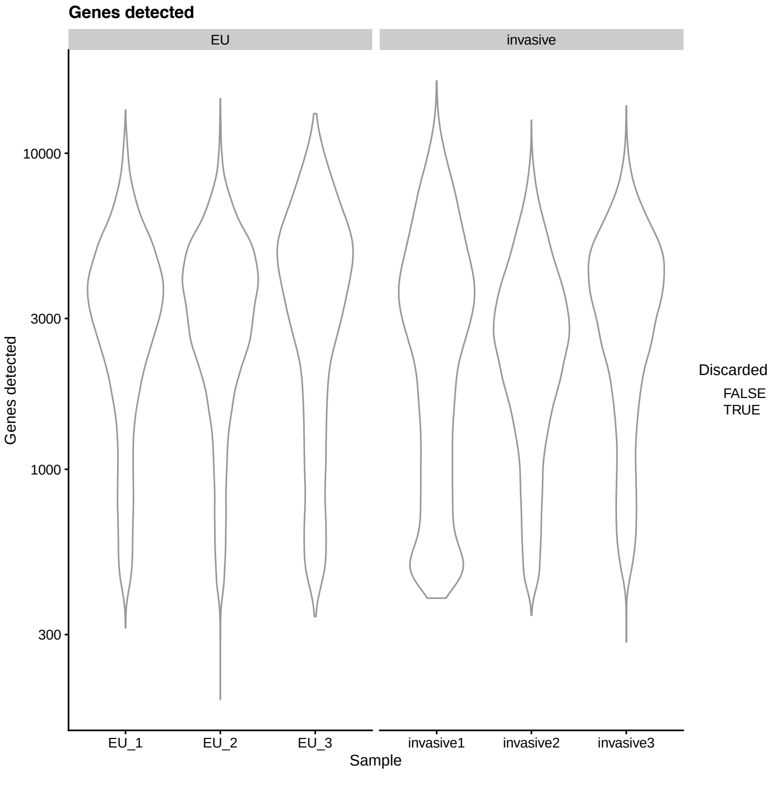

**Supplementary Figure S3 Quality control of single-cell RNA-seq data: chloroplast and mitochondrial content across EU and invasive samples.** Cells with more than 30% chloroplast transcripts (left panel) and more than 10% mitochondrial transcripts (right panel) are highlighted in blue, indicating potential low-quality or stressed cells that may be excluded from downstream analysis to improve data quality.

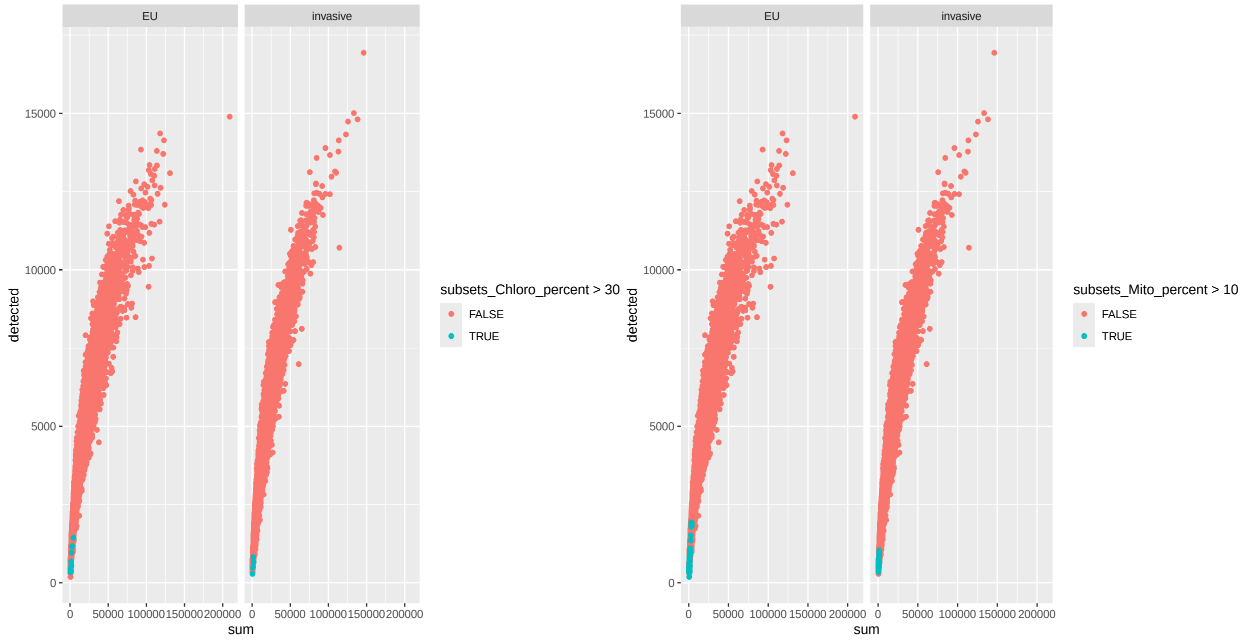


**Supplementary Figure S4** The t-SNE projection of single-cell transcriptomic data clustered at a resolution of 0.8. Each dot represents an individual cell, colored by its assigned Seurat cluster (0–18).

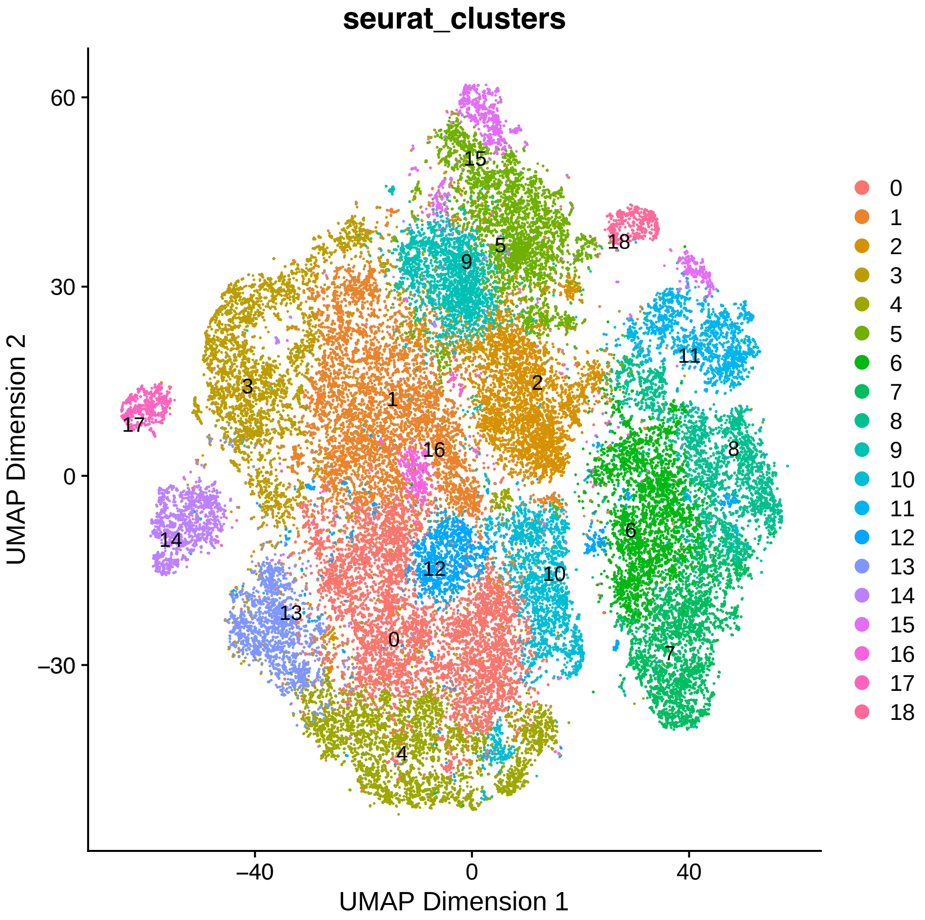


**Supplementary Figure S5** UMAP projection of single-cell transcriptomic data, split by sample identity. Each panel represents a separate sample, with cells colored according to their respective UMAP coordinates. The samples are labeled on the x-axis, and the coordinates reflect the dimensionality reduction based on the top principal components of the data. The plots are arranged in a 2-column layout to facilitate comparison of UMAP embeddings across samples.

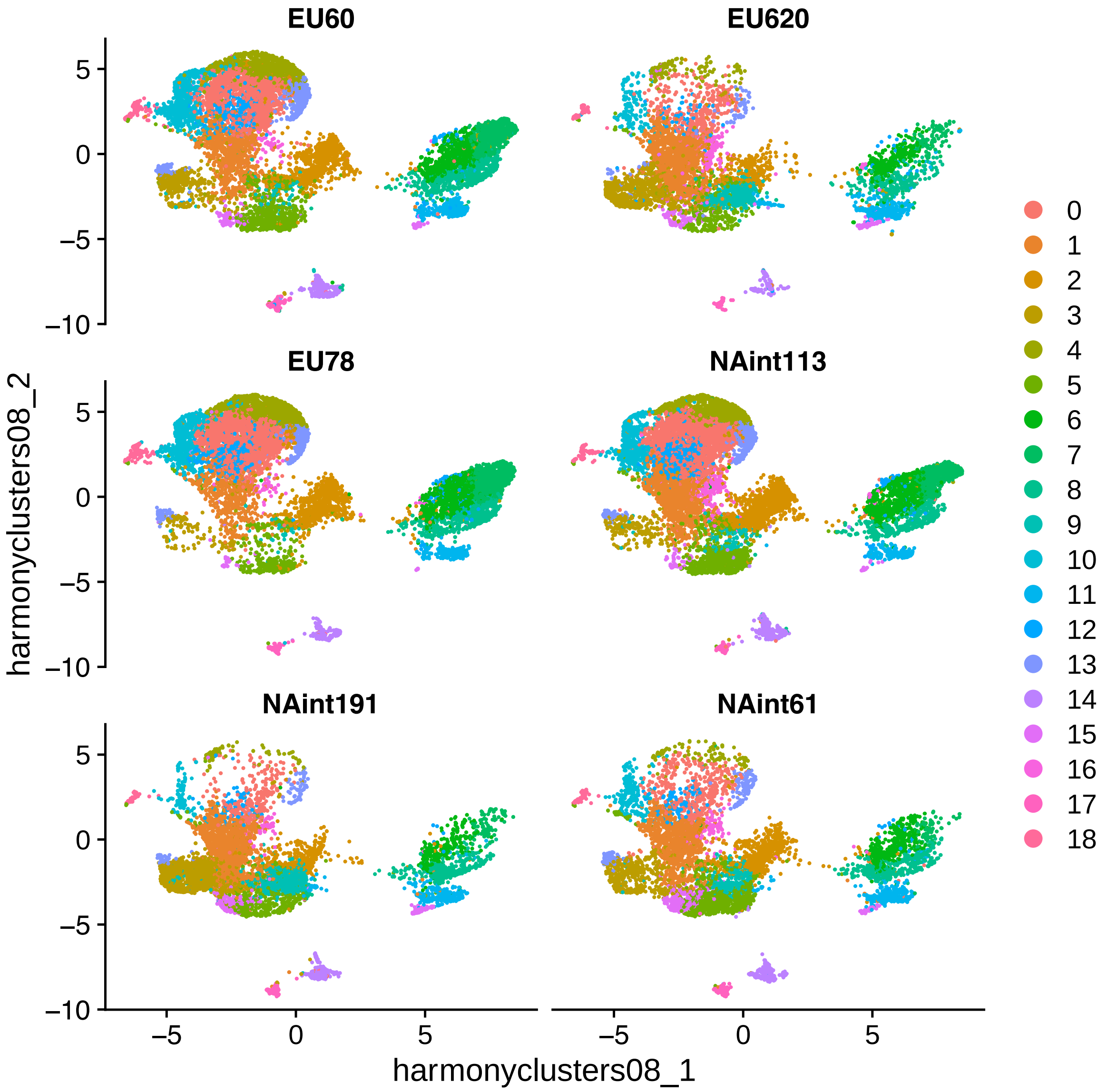


**Supplementary Figure S6** Heatmap of gene expression levels in spatial transcriptomic tissues. Spatial transcriptomic data is overlaid on histological tissue sections, with each spot representing a spatial capture location. The color gradient indicates the number of detected genes (nGene), ranging from low (blue) to high (red) expression.

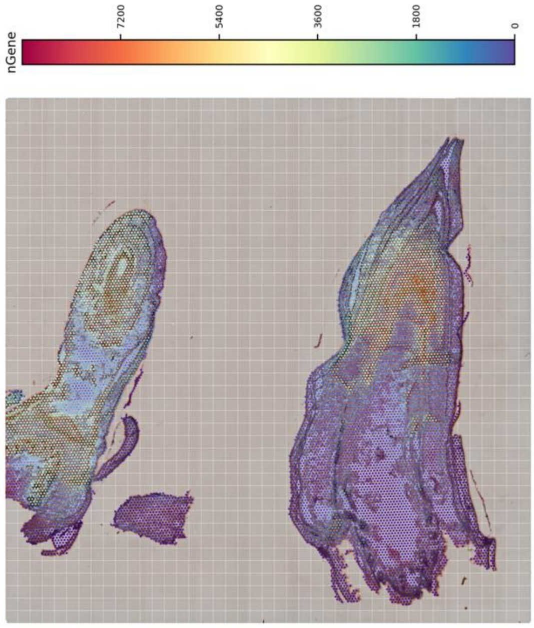


**Supplementary Figure S7** Nineteen clusters identified from spatial transcriptomics data at a spatial resolution of 37 μm, using level 7 subspot granularity. This clustering reveals the spatial organization of transcriptionally distinct regions within the tissue.

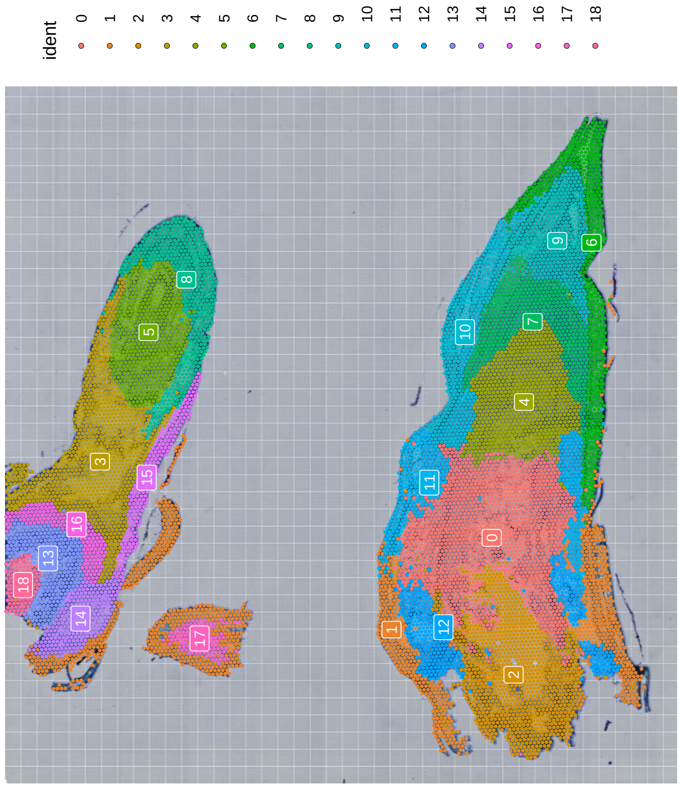

**Supplementary Figure S8** Spatial projection of single-cell transcriptomic clusters onto the spatial transcriptomics dataset. Each mapped cell cluster reflects its inferred spatial localization within the tissue, enabling integration of single-cell resolution with spatial context.

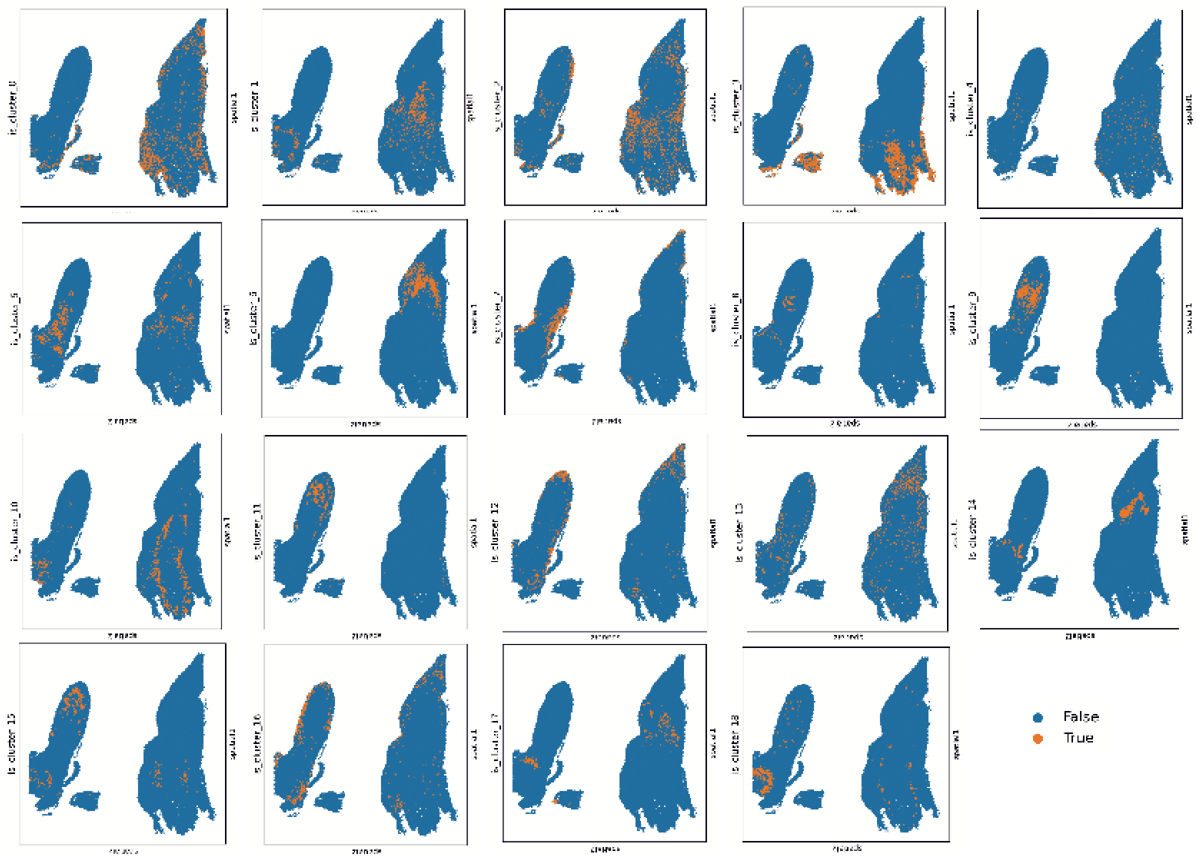


**Supplementary Figure S9** Significantly enriched Gene Ontology (GO) terms identified for each cluster. These annotations highlight the predominant biological processes associated with the gene expression profiles of each cluster.

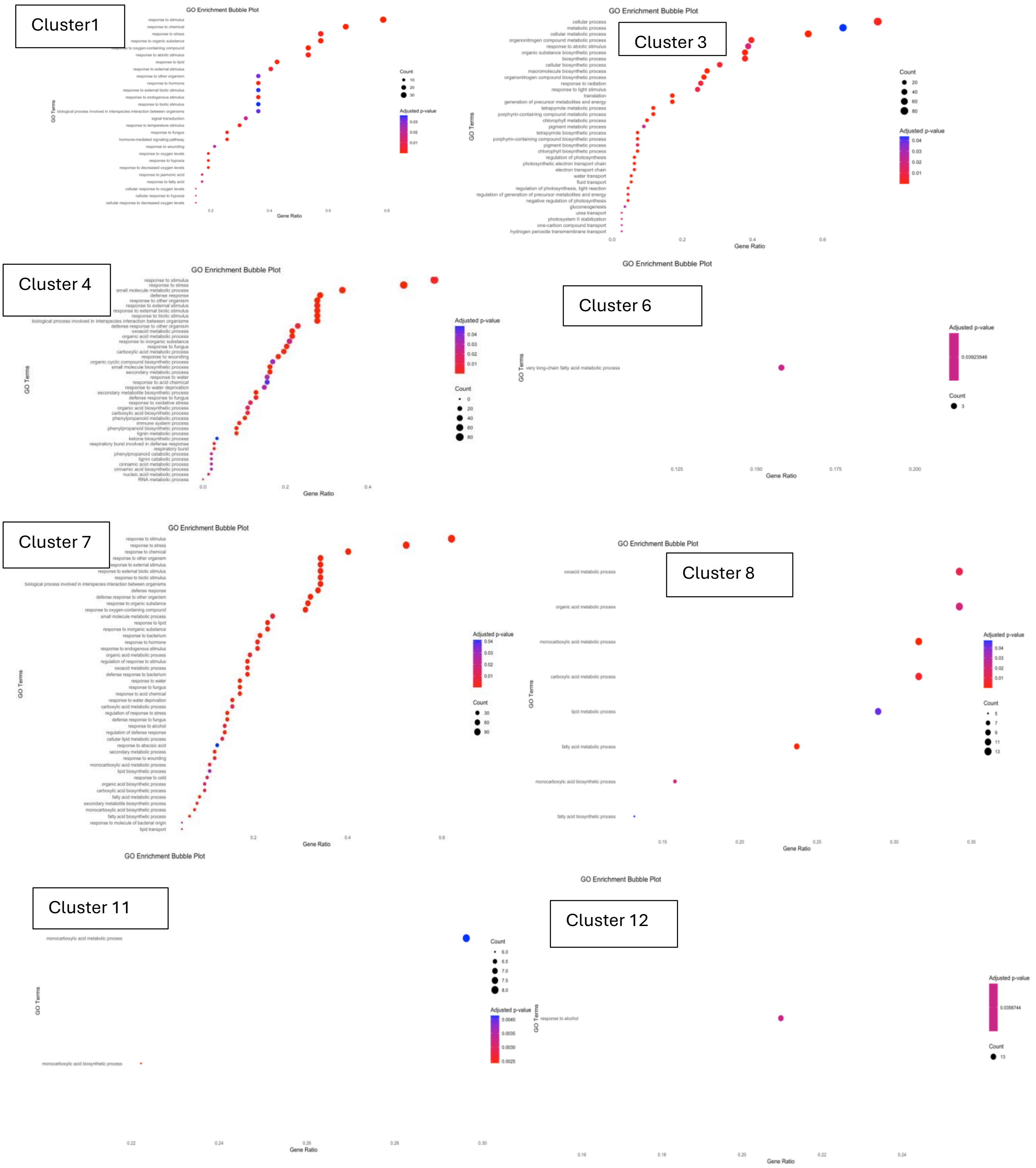
Cluster1
GO Enrichment Bubble Plot
Cluster 3
GO Enrichment Bubble Plot
Cluster 4
GO Enrichment Bubble Plot
Cluster 6
GO Enrichment Bubble Plot
Cluster 7
GO Enrichment Bubble Plot
Cluster 8
GO Enrichment Bubble Plot
Cluster 11
GO Enrichment Bubble Plot
Cluster 12
GO Enrichment Bubble Plot
GO Terms
Gene Ratio
Count
Adjusted p-value

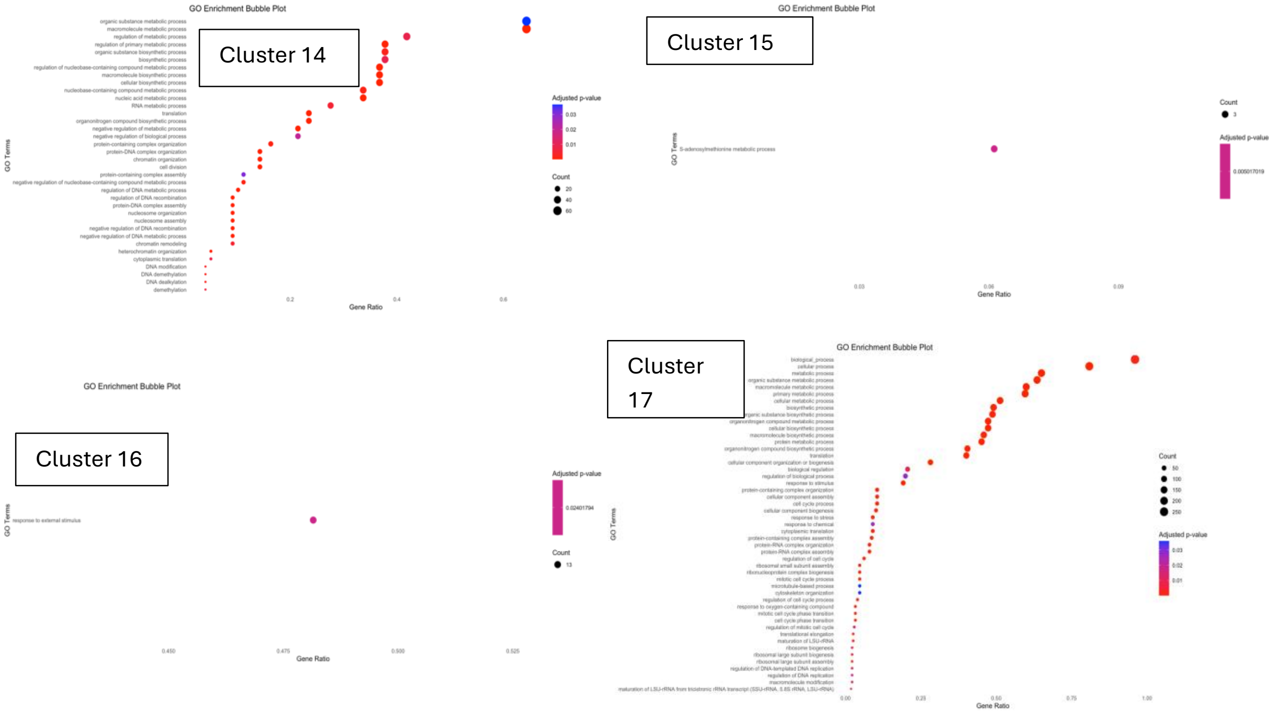


**Supplementary Figure S10** Inferred fate probability of  sample EU620 following dpt pseuodotime trajectory.

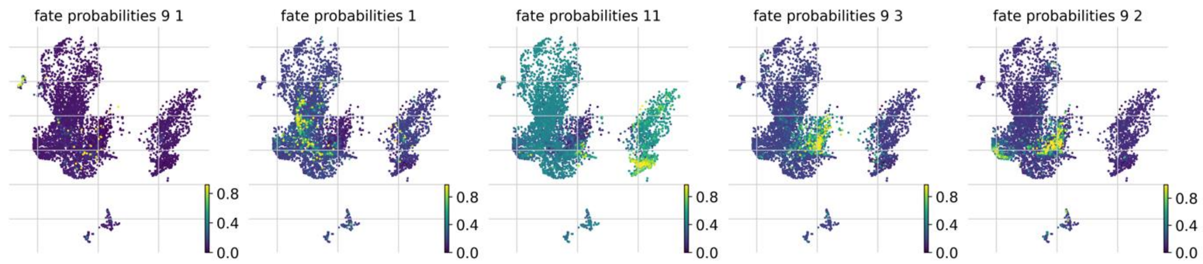


**Supplementary Figure S11** Driver genes and gene expression levels across different states of Cluster 9 along the pseudotime trajectory. Panel a,b, c shows cell state 1, 2, 3 respectively.

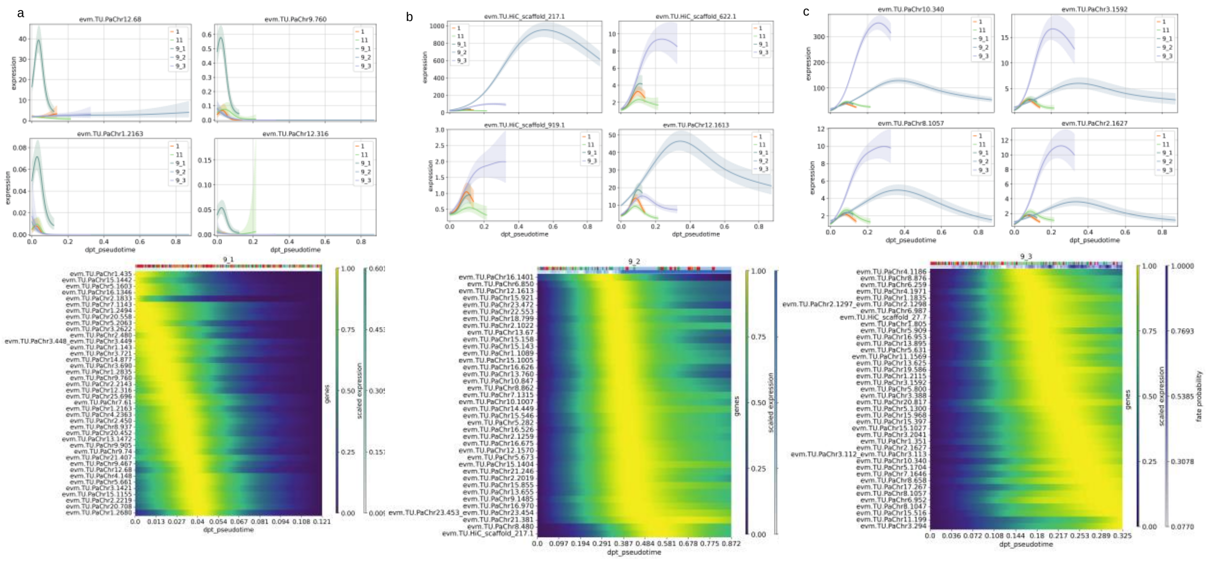

**Supplementary Figure S12** Number of cells per cluster across individual samples. Samples EU60, EU620, and EU78 represent non-invasive European populations, whereas NAint113, NAint119, and NAint61 correspond to invasive North American populations.

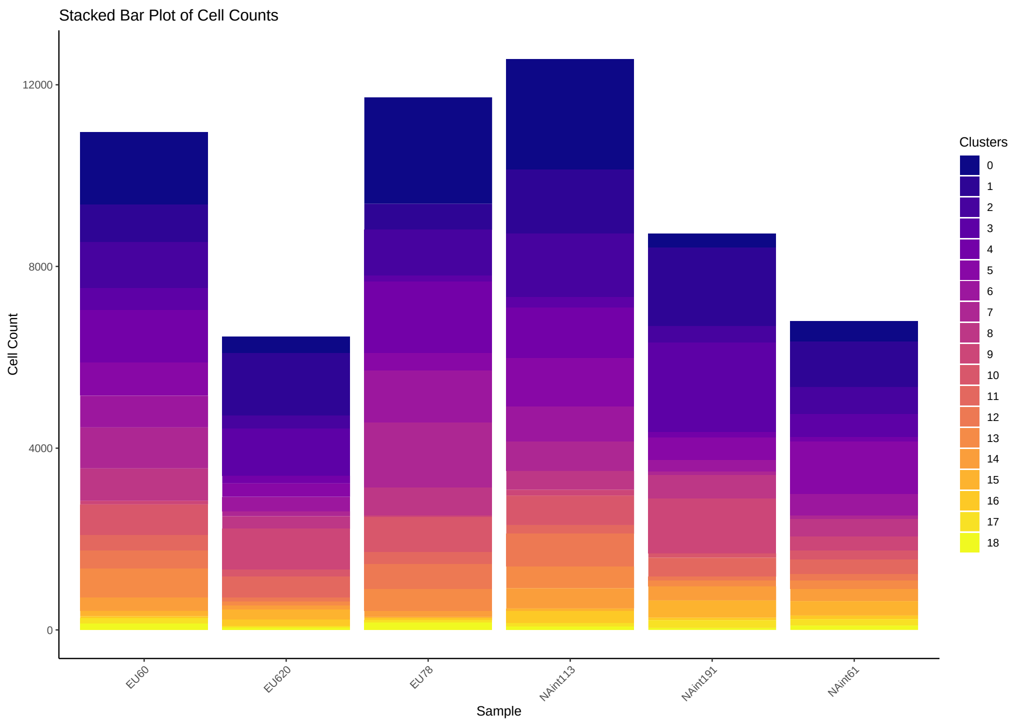


**Supplementary Figure S13** Visualization of the first two principal components (PCA) based on pseudobulk expression profiles from each cluster and all clusters in the single cell dataset across each sample. For each cluster, the invasive and ancetral European populations were seperated on PC2, and a large proportion of the variation was detected in PC1, which represents the gene expression variations within each population.

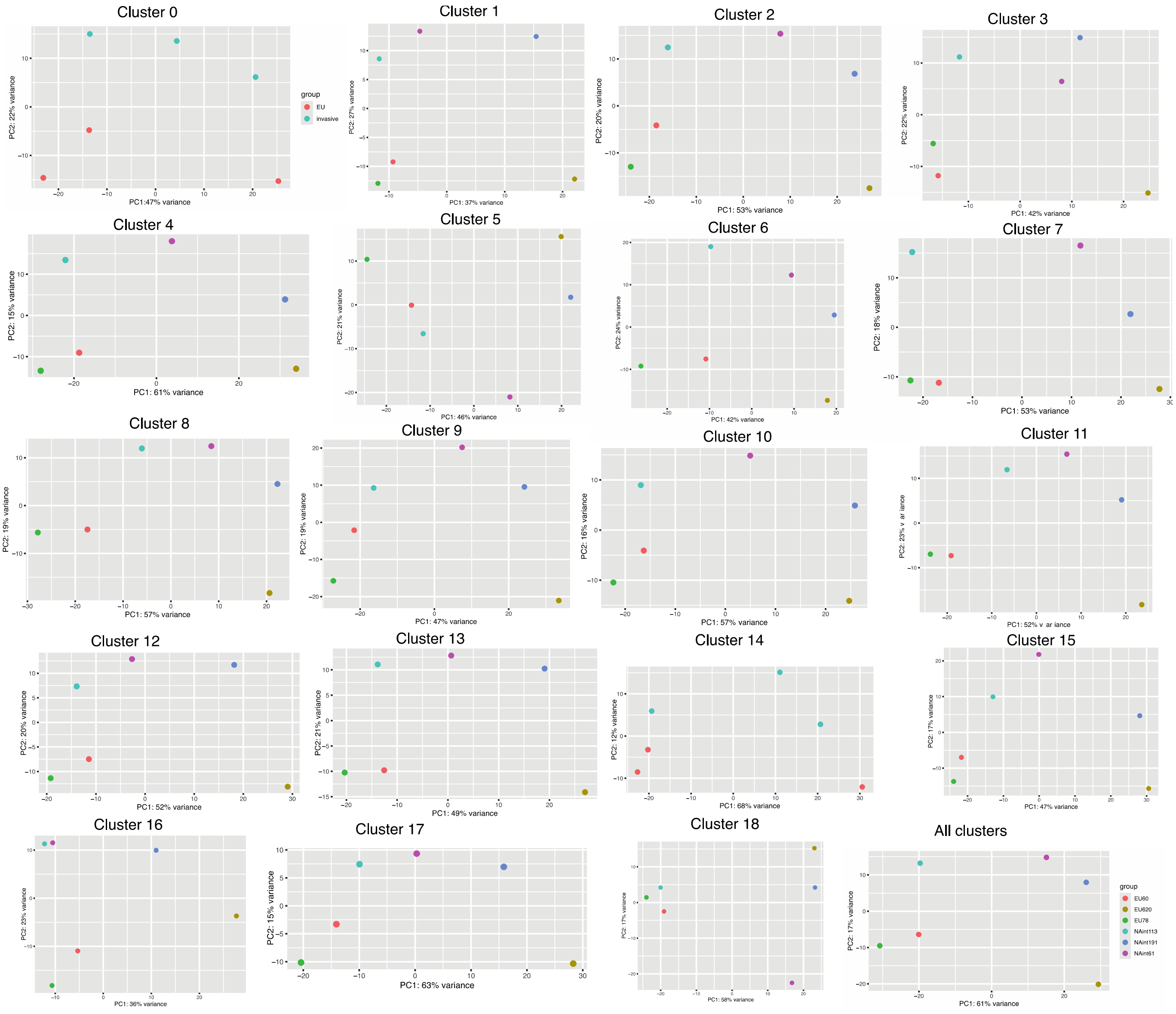


**Supplementary Figure S14** RNA velocity analysis for each sample, depicting transient transcriptional dynamics within the shoot system. Arrows represent the inferred direction and magnitude of RNA flow, indicating the potential future states of individual cells and highlighting developmental trajectories across distinct tissue types.

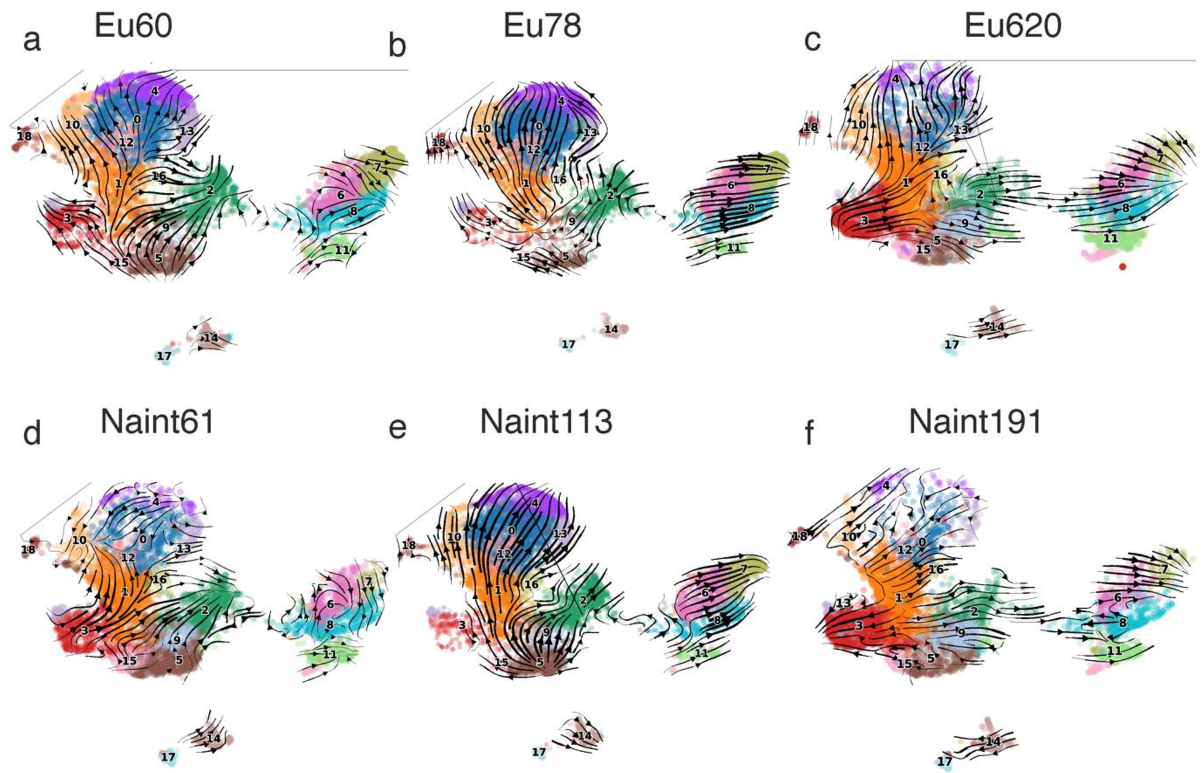


**Supplementary Figure S15 heatmap of sample correlation from cluster 0 to 18.**

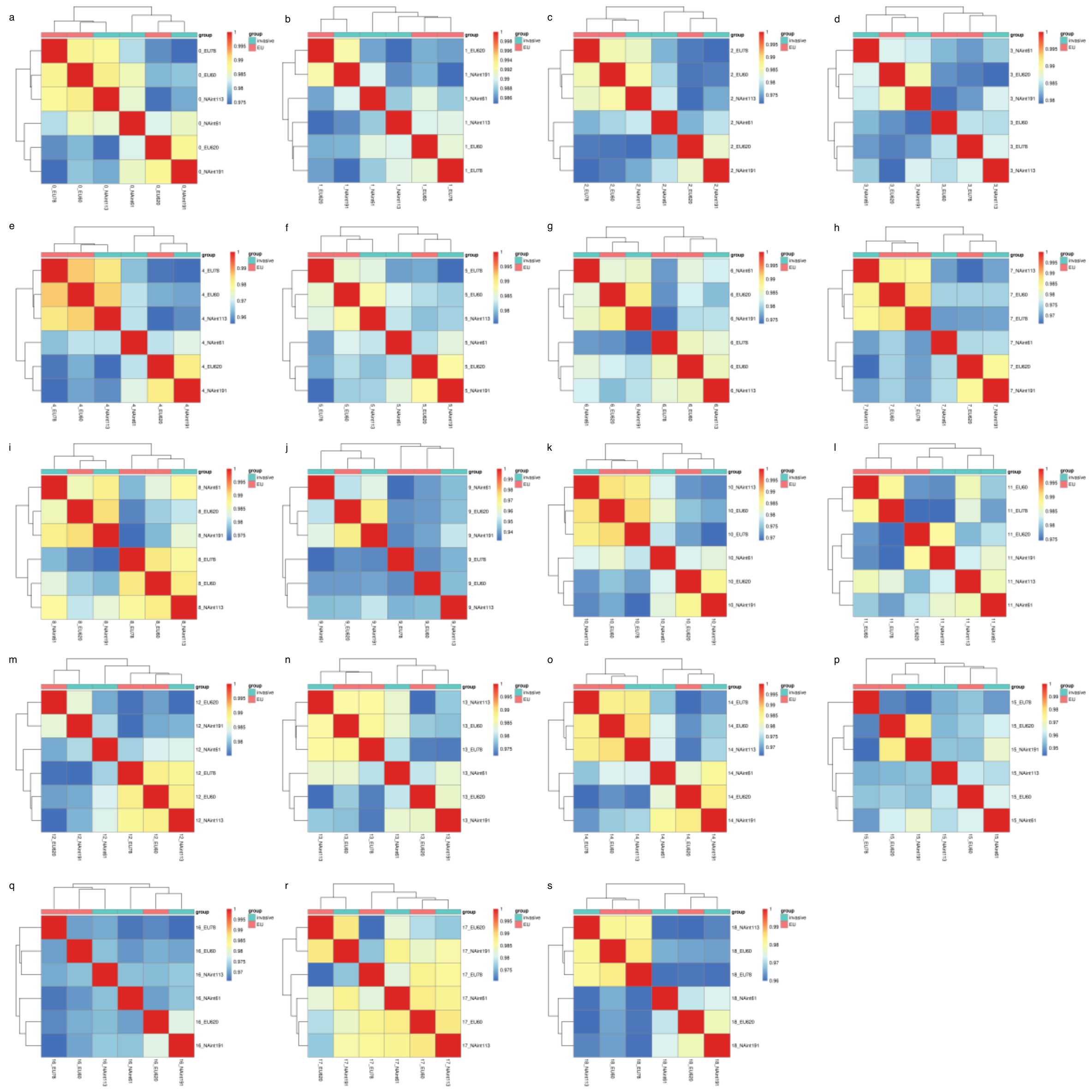

**Supplementary Figure S16. Fst values between the EU and invasive populations along the genome, using a sliding window size of 10 kb.**

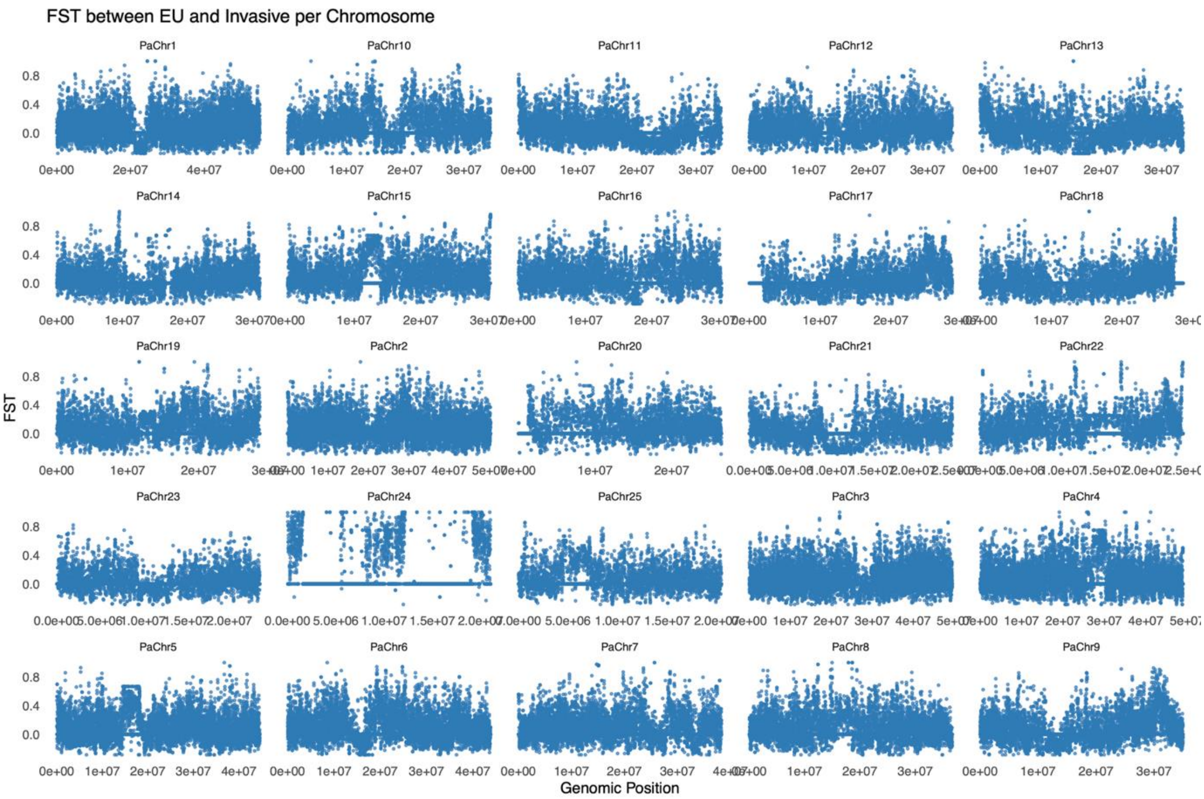


**Supplementary Figure S17. Tajima's D values for the invasive and EU populations on each chromosome, estimated using a sliding window size of 10 kb.**

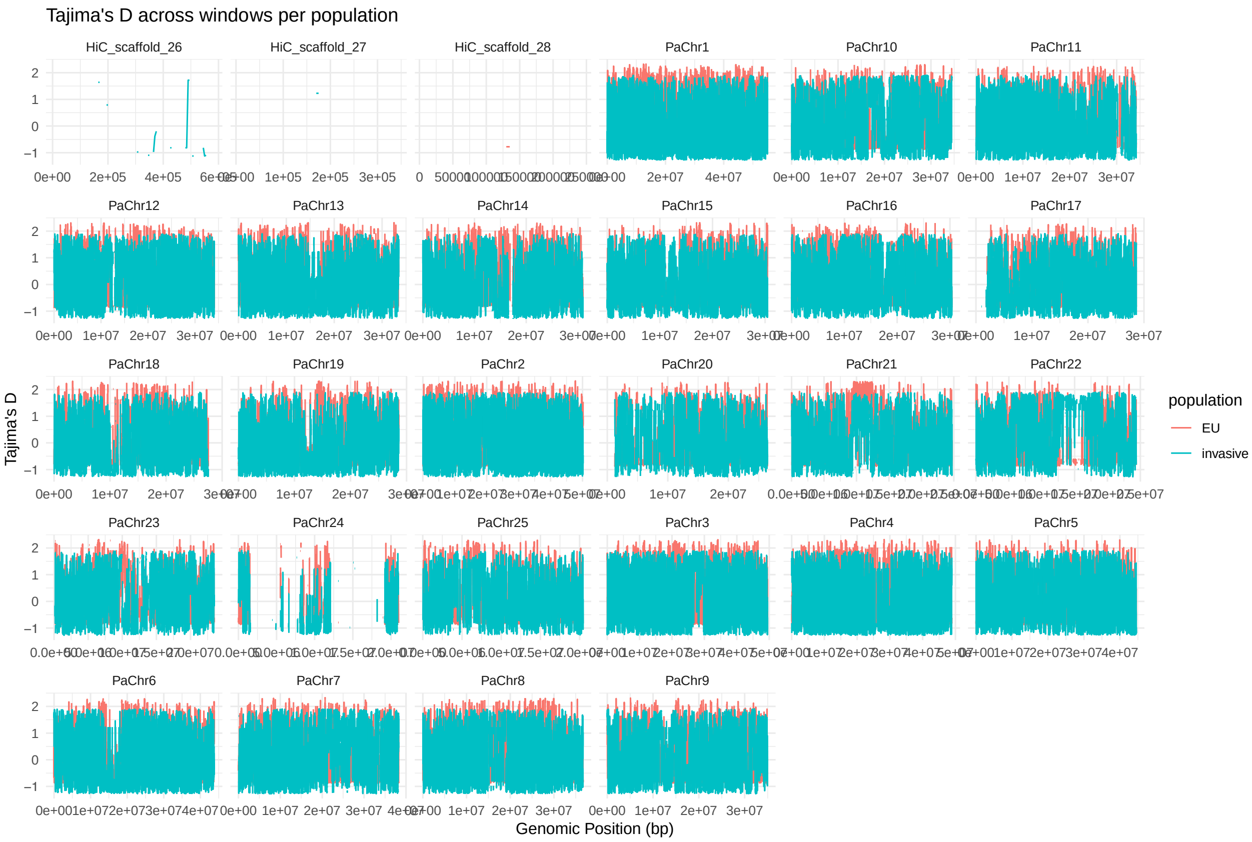

**Supplementary Figure S18 Phylogeny of two additional genes from the B chromosome, Illustrating the origin of B chromosome genes.**

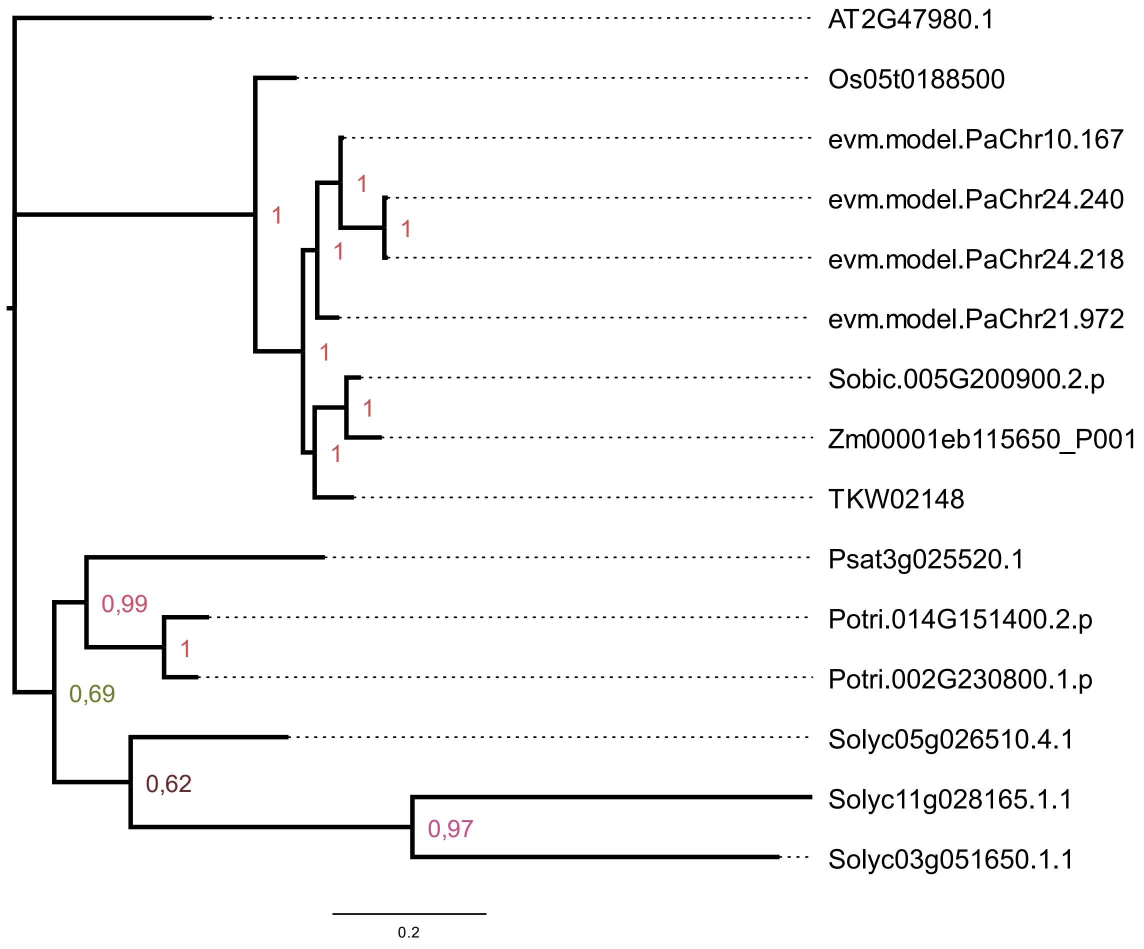


### Supplementary Note 1

Using two replicates of single cell dataset, for the SAM cluster, among the 442 DEGs, 279 genes were upregulated in the invasive lineage, while 163 genes were downregulated. The upregulated genes were primarily enriched in processes related to light harvesting in photosynthesis, translation, chlorophyll biosynthesis, response to abiotic stimuli (including radiation, cold, high and low light intensity), photosynthetic electron transport chain, plastid translation, granum assembly, and membrane bending. On the other hand, the downregulated genes were enriched in processes related to stress response, defense response, and lipid transport.

### Supplementary Note 2

One gene, SCC3, located on the B chromosome, was consistently upregulated across all three clusters. In Cluster 14, most of the upregulated DEGs are primarily involved in photosystem functioning (PSAG, PSAH2, PSAN, PSBP1, LHCB6, PSBW, LHCB4.2, PSBQA, LHB1B2), chloroplast accumulation (CPN60A), chloroplast protein translation (PRPL34), and plant hormone response (XTH24, PYL11). In Cluster 11 (epidermal cell), the top upregulated genes are primarily involved in anthocyanin biosynthesis (ANS, CHS, DFR). In Cluster 18, the upregulated genes encode proteins involved in environmental stimuli responses (TCH4, SUS4, PYL11) .
